# Every Cure Knowledge Graph: A Unified Biomedical Knowledge Graph for Drug Repurposing

**DOI:** 10.64898/2026.08.26.747253

**Authors:** Piotr Kaniewski, E. Kathleen Carter, Dan Rhodes, En May Lim, Jane Li, Jacques Vergine, Nico Matentzoglu, Kevin Schaper, Jason Reilly, Shilpa Sundar, Laurens Vijnck, Elliot Sharp, Nelson Alfonso, Amy Ford, Alexei Stepanenko, Charlie Hempstead, Pascal Brokmeier, Christopher Bizon, Alexander Tropsha, Melissa A. Haendel, David C. Fajgenbaum, Lee Lancashire

## Abstract

Identifying causal connections between existing drugs and mechanistic profiles of diseases is a foundational step for effective drug repurposing. Although knowledge graphs (KGs) are highly suited for consolidating biomedical databases and tracking these connections, a single biomedical KG is constrained by its ingestion pipeline and knowledge sources. While different biomedical KGs could be complementary if combined, efforts to combine them into a unified and more comprehensive KG are hindered by lack of interoperability and poor provenance. To address those issues, we present EC-KG, a Biolink Model-compatible KG for computational drug repurposing. EC-KG is an interoperable, provenance-first KG which integrates RTX-KG2, ROBOKOP, and PrimeKG at the network-level, encapsulating over 7 million nodes and 81 million edges from 95 primary data sources. EC-KG has improved coverage of core biomedical entities such as drugs, targets, and diseases relevant to drug repurposing vs source graphs, and captures complex biomedical mechanisms within its topology. We demonstrate that the network unification in EC-KG leads to emergence of novel, mechanistically relevant pathways which are disconnected in the underlying constituent networks and show its applications in method development, benchmarking and predictive drug repurposing applications. EC-KG has already been successfully used in drug repurposing research to surface Botulinum Toxin A as a candidate to treat Major Depressive Disorder, as well as to validate repurposing of Lenalidomide and Dexamethasone for a subgroup of patients with Rosai-Dorfman Disease.

## Background and Summary

Drug repurposing is the process of identifying new therapeutic applications for existing medicines.^1,2^ Much like traditional drug discovery, drug repurposing leverages the mechanistic understanding of drug and disease targets to identify therapeutic candidates. However instead of characterizing a new compound and its efficacy profile *de novo*, it seeks to discover a relationship between targets of an approved drug with mechanisms in a disease beyond the drug’s established indications.^1,3^ Several well-known examples highlight its potential, including the repurposing of thalidomide for multiple myeloma after initially being used for leprosy and rituximab for rheumatoid arthritis after initially being used for lymphoma.^4–6^

While drug companies often successfully repurpose drugs during the early stages of their patent lives, serendipitous observations by clinicians and researchers have historically been the main path to successful drug repurposing after drugs lose patent exclusivity.^6–8^ However, the rapid expansion of biomedical data has made systematic approaches to identifying repurposing opportunities possible at scale. Biomedical knowledge graphs (KGs), with their ability to model complex and large-scale data are well suited to the querying, reasoning and inference tasks necessary for drug repurposing. The node and edge based structure of a KG is naturally suited to representing biological systems such as gene and protein interaction networks, phenotypic associations and biological processes. KGs integrate data from disparate sources into a standardized, machine-readable representation, enabling the application of rule- or ML-based link prediction models.^3,9–14^ Examples include the Monarch KG, used for rare disease diagnostics;^15^ PrimeKG, used as a network for the training of foundational models such as TxGNN or NetMedGPT^16–18^; RTX-KG2, used for the development of the KGML-xDTD system, which successfully identified a TNF inhibitor that was effective for treating a patient with idiopathic Multicentric Castleman Disease.^19–21^; and SPOKE and ROBOKOP KG, used for development of disease-specific algorithms for repurposing against diabetes and COVID-19 respectively. ^22–26^ These knowledge graph based approaches served as the initial methodologies for advancing the novel approach of computational pharmacophenomics, which involves systematic quantitative evaluation of the potential of all drugs (pharmaco-) as treatments against all diseases and even sub-phenotypes within a disease (-phenomics).^27^

Despite their demonstrated usefulness for drug repurposing and computational pharmacophenomics, current biomedical KGs face three main systematic limitations, the first of which is completeness.^28–30^ A biomedical KG is inherently restricted by what the world knows about biology. There is both missing information that the world hasn’t discovered as of yet, such as an effective therapeutic target in the vast majority of amyotrophic lateral sclerosis patients, and incorrect information that is propagated in the medical literature. Beyond these global challenges that are very difficult to address, there are related challenges around which datasets should be selected as primary sources and its preprocessing strategy.^30^ For example, entities such as *Azacitidine*, *CDKN1B* protein, and *Multiple Endocrine Neoplasia Type 4,* are captured in both RTX-KG2 and ROBOKOP KG however relationships connecting the entities are inconsistently present across the networks: drug-protein edge only exists in ROBOKOP KG while protein-disease edge only exists in RTX-KG2. This has critical implications on discovery of novel therapeutics as it may lead to the overlooking of causal, mechanistic pathways which are not captured within a set of ingested databases. Beyond asymmetrically captured relationships, KGs can have substantially different entities representing the same system. For example, ROBOKOP KG and PrimeKG ingest Monarch’s Mondo Disease ontology (Mondo) to build disease and phenotype networks, but the latter groups clinically similar diseases into a single entity, driven by a greater clinical focus. This results in fewer disease entities (22,236 vs 17,080) in PrimeKG.^16^ While clinically reasonable, excessive merging leads to information loss - for instance both MONDO:0014036 and MONDO:0014265 represent Alzheimer disease but caused by mutation of *TREM2* and *ADAM10* genes respectively. Such lack of coverage of biomedical entities can have deleterious implications for computational repurposing, which often relies on genetic detail.^27^

The second systematic limitation is the lack of standardization and normalization, which leads to challenges in interoperability.^31–33^ Tools such as Monarch’s Biolink Model help in the standardization of KGs by defining their data model, node, and edge schemas.^34, 39^ While such tools are useful, they do not address normalization, the process of resolving multiple identifiers for a single biomedical concept into a unified and consistent representation. For example, across various KGs and databases, the concepts “water” and “aspirin” are represented by over 30 and 400 different compact URIs (CURIE) respectively.^35,36^ Normalization resolves these, preventing duplication of information and possible fragmentation of the graph. The proper handling of both standardization and normalization, allows for graphs to be interoperable. Interoperability is essential for the seamless exchange and comparison of information between data sources. While FAIR data frameworks detail how to best achieve this, many biomedical KGs prioritize use-case specific utility over alignment with FAIR principles.^37^ For example, despite RTX-KG2 and ROBOKOP KG both being Biolink compatible, the different normalization frameworks used for both (ARAX^38^ and Babel^32^ respectively) results in a lack of interoperability, as we cannot reliably know that “water” in RTX-KG2 is the same biomedical concept as in ROBOKOP KG.^36^ This creates barriers to important aspects of research such as evaluation of machine learning (ML) models, as the lack of a single network and data model restricts the ability to benchmark approaches by comparing like for like.^40^ Therefore, lack of interoperability hinders comparative evaluation of computational methods and prevents integration of complementary data from independent KGs.

The final limitation is provenance-tracking. Provenance is essential to establish the relative importance and quality of nodes and edges as KGs integrate heterogenous data sources. While provenance can in principle be traced through a KG’s construction pipeline, the data is often missing from the edge or node level attributes in the graph.^33^ Furthermore, data may have been through several transformations, such as in the case where biomedical databases used in a KG aggregate the same source data using different techniques. Such instances require in-depth provenance, tracking metadata such as knowledge extraction methods, or publications supporting a specific relationship. For example, the Nicotinic Acid-HCAR2 relationship is encoded in both RTX-KG2 and ROBOKOP. The relationship is supported by 19 publications within SemMedDB^41^ (via RTX-KG2) and 10 publications with quantitative descriptors in Pharos^42^ (via ROBOKOP). Currently, biomedical KGs vary considerably in how well they track these essential attributes, leaving edges poorly characterized and reducing confidence of drug repurposing predictions for both researchers and computational agents.

The issues of completeness, interoperability, and poor provenance are well-recognized challenges across the biomedical KG field.^33,43^ While several efforts have attempted to address these challenges, including integration frameworks such as Petagraph^44^ and DINGO,^45^ or unified biomedical KGs such as DRKG^46^ and SPOKE^22^, their primary method of adding information to the network is through addition of new source databases, requiring a complete reconstruction of the network to add new knowledge, or enrichment of the network with non-KG data like omics through manual ontology mappings, requiring significant time and human effort. None of the frameworks have achieved a reusable KG-level integration of multiple biomedical networks while retaining an interoperable backbone with both source- and graph-level provenance.^33^

In this work, we present Every Cure Knowledge Graph (EC-KG), an integrated knowledge graph designed for drug repurposing, explicitly addressing the challenges of completeness, interoperability, and provenance that limit existing biomedical KGs. EC-KG is constructed through a cloud-agnostic and open-source framework that provides infrastructure for the standardization and harmonization of biomedical knowledge graphs (Fig. 1). Through preprocessing and unification of PrimeKG, RTX-KG2, and ROBOKOP, EC-KG forms a Biolink Model compatible network comprising over 7 million nodes and 81 million edges, with more than 60 unique node types and 91 unique edge types. By aggregating 95 primary knowledge sources, EC-KG provides a rich landscape for drug repurposing and discovery. EC-KG captures complementary information from existing biomedical networks and enables the identification of novel mechanistic paths supporting computational discovery that are often inaccessible within any single biomedical network (Fig. 1d). Additionally, unification of the metadata layer makes EC-KG edges information-rich, strengthening the evidence supporting the relationship and increasing the confidence of drug repurposing predictions, both for researchers and computational agents.

**Figure 1.**
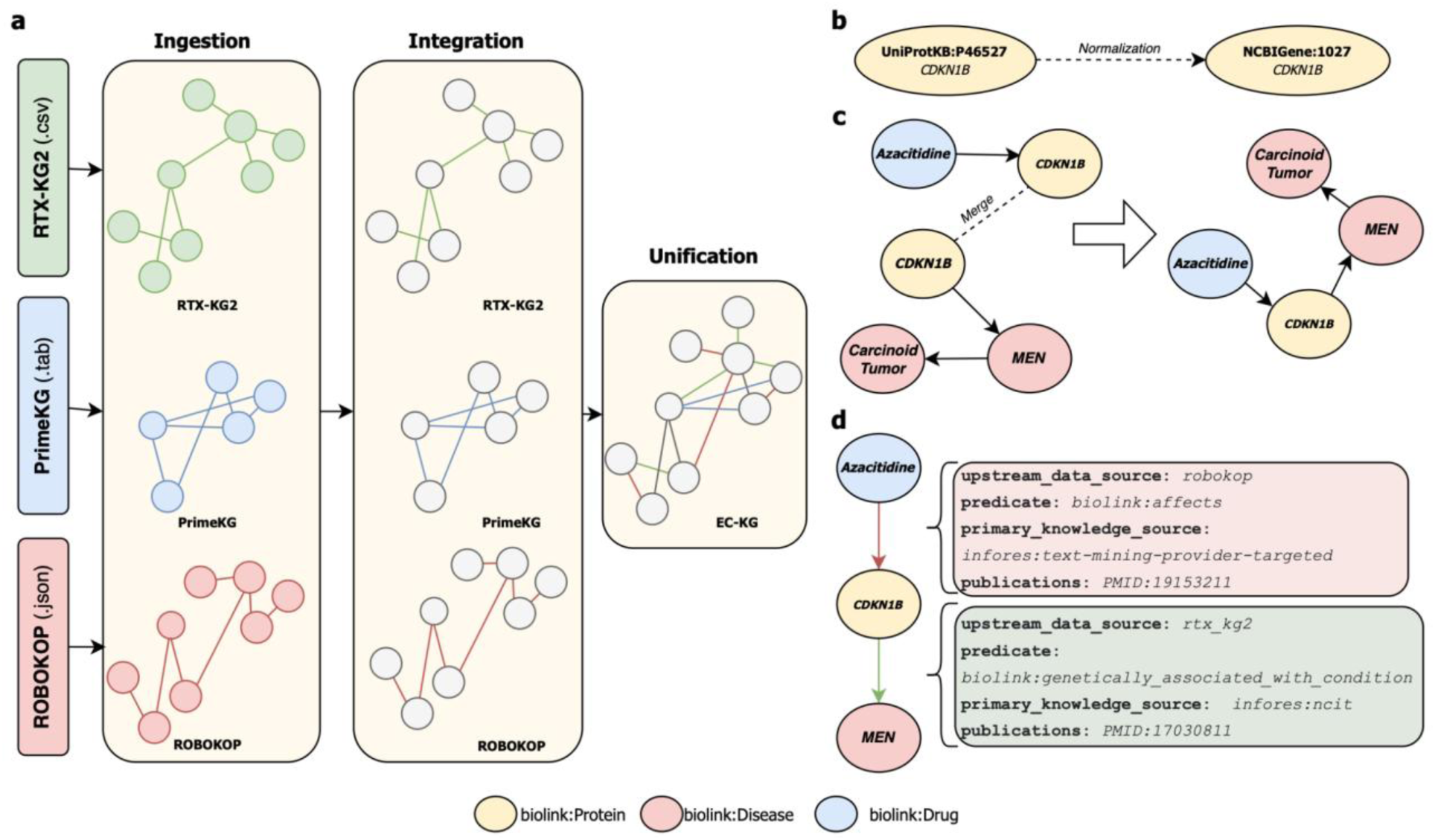
Schematic representation of EC-KG construction pipeline. a) EC-KG is constructed by ingesting, preprocessing, integrating, and unifying three different biomedical KGs into one interoperable biomedical network. b) Node normalization process ensures entities and their identifiers across the upstream KGs are resolved to a consistent biomedical vocabulary. c) The unification process joins separate knowledge graphs into a harmonized network by merging identical entities across graphs. d) Resulting subgraph extracted from EC-KG, showing multi-level provenance of upstream KGs and primary knowledge sources.

Public biomedical KGs vary considerably in scale and design. Among Biolink Model-compatible networks, RTX-KG2 is the largest by node and edge count (6,763,921 nodes; 46,366,733 edges; 64 knowledge sources), followed by ROBOKOP KG (4,663,956 nodes; 26,268,290 edges; 45 knowledge sources) and MonarchKG (1,128,873 nodes; 12,154,597 edges; 87 knowledge sources), all of which utilize different node resolution frameworks such as ARAX or Babel. Non-Biolink KGs are comparatively smaller: PrimeKG (129,375 nodes; 4,050,249 edges; 20 sources), DRKG (97,238 nodes; 5,874,261 edges; 6 knowledge sources), Hetionet (47,031 nodes; 2,250,197 edges; 29 knowledge sources), and PharmKG (188,296 nodes; 1,093,236 edges; 7 knowledge sources) all employ custom, non-standardized data models and normalization frameworks. This heterogeneity in scale, source coverage, and standardization reflects a broader, community-recognized challenge in assessing and comparing biomedical KG quality.^33^ EC-KG substantially exceeds all of these individually, with 7,354,612 nodes, 81,149,812 edges, and 95 aggregated knowledge sources, while enforcing Biolink Model compatibility and Babel-based normalization.

## Methods

A schematic representation of the EC-KG construction pipeline is shown in Figure 1. The pipeline is implemented using the Kedro^49^ framework and consists of three modular stages: preprocessing, ingestion, and integration. Each stage contains a set of transformations for processing of each biomedical KG and their integration into EC-KG. Raw, intermediate and final data products, including quantitative metrics, are stored as parquet files according to Kedro’s layered data architecture. Each stage of the pipeline, along with the FAIR-design and infrastructure supporting data intensive processing, is described in the following sections.

### Core drugs and diseases for repurposing

To ensure EC-KG meets the needs of drug repurposing research, we defined a core set of drugs and diseases which we consider to be most relevant for repurposing, as well as the relationships between them, which we use throughout to assess EC-KG and other graphs.

We manually compiled the core drug list (N=1,784) from FDA- and EMA-approved drugs, with closely related drug forms collapsed into one entity, such as different salt forms of a drug or drugs and their prodrugs.^50^ For example, Warfarin potassium and Warfarin sodium are consolidated as warfarin, and Amiloride hydrochloride is consolidated with Amiloride. To compile our core disease list, we started by extracting a disease list from the MONDO ontology.^51^ The Mondo ontology contains a hierarchy of over 22,236 entities including high-level disease groupings which are not specific to one clinical condition, such as ‘nervous system disorders’. Additionally, it contains numerous clinically identical subtypes of disease, such as over 100 genetic subtypes of Retinitis pigmentosa. We found that repurposing opportunities made on these highly specific conditions were significantly less actionable as repurposing predictions, and so focused on conditions widely recognised by clinicians and patients. To identify these, an internal physician manually annotated Mondo diseases based on their clinical suitability for drug repurposing, yielding a shortened core disease list (N=7,270).

In addition to drug and disease lists, we manually curated a list of drug-disease pairs congruent with the core drug list and disease list representing on-label indications (N=2,034) and off-label (N=746) uses. Off-label uses are particularly important for drug repurposing as they highlight clinically successful applications of drugs in conditions outside their approved indications. We constructed the dataset through the manual inspection of sources such as clinical guidelines, DailyMed and Drug Central, including only interventional treatments and excluding pharmaceutical interventions used for prophylaxis (e.g., vaccines for infectious diseases), diagnosis (e.g., allergens for allergy testing) and symptomatic treatment of diseases (e.g., analgesics for painful conditions). Each curator made independent curation decisions; in cases of uncertainty, a second reviewer was consulted to reach consensus. This list of on-and off-label drug-disease relationships was used for technical validation of EC-KG, and it can be also leveraged as a high-quality evaluation dataset for downstream applications. Importantly, we constructed the core drug, disease, and indications lists independently of EC-KG’s source data. Of the 2,780 indication edges identified, 2,418 can be mapped to edges in EC-KG.

### Data Unification

EC-KG was constructed through three distinct stages: preprocessing, ingestion, and integration.

#### Preprocessing

As biomedical KGs vary in their data models, standardization frameworks, and attributes, harmonization of the networks into a semantically consistent format is required (Table 1). Therefore, we individually processed each upstream KG to ensure all biomedical networks are aligned with the Biolink Model to ensure semantic consistency. Furthermore, we distributed processed networks in KGX format to facilitate knowledge graph exchange through standardized format structure by using identifiers in CURIE format.^52^

**Table 1.** KGs ingested in the raw format. Identifier format specifies the canonical standard of primary identifier attributes. Both ROBOKOP KG and RTX-KG2 largely derive their attributes fromBiolink Model. PrimeKG has 4 edge attributes and 5 node attributes however drug and disease nodes contain 17 and 14 features respectively.

| KG | Size | Data Model | KGX format | Type | Normalization | Attributes Number |
| --- | --- | --- | --- | --- | --- | --- |
| RTX-KG2 | 6 763 921 nodes<br>46 366 733 edges | Biolink | True | 56 node types<br>69 edge types | ARAX | 15 edge attributes<br>10 node attributes |
| PrimeKG | 129 375 nodes<br>4 050 249 edges | PrimeKG Specific | False | 10 node types<br>30 edge types | Custom | 4 edge attributes<br>5 node attributes |
| ROBOKOP KG | 4 663 956 nodes<br>26 268 290 edges | Biolink | True | 79 node types<br>68 edge types | Babel | 66 edge attributes<br>35 node attributes |

**Table 2.** Coverage of drugs and diseases across upstream knowledge graphs and EC-KG with core drug and disease list connectivity. The metrics indicate minimal fragmentation of the network. LCC - Largest Connected Component; WCS - Weighted Connectivity Score. Drugs missing from LCC are due to upstream RTX KG2 containing floating entities, which then get removed upon KG unification.

|  | Drug List | Disease List |
| --- | --- | --- |
| **Total Count** | 1,784 | 7,270 |
| **PrimeKG Coverage** | 1,392 (78%) | 6,640 (92%) |
| **ROBOKOP KG Coverage** | 1,655 (92%) | 7,191 (99%) |
| **RTX-KG2 Coverage** | 1,720 (95%) | 7,165 (99%) |
| **EC-KG Coverage** | 1,762 (99%) | 7,193 (99%) |
| **Core Drugs and Diseases in LCC** | 1,762 | 7,193 |
| **LCC Fraction** | 1 | 1 |
| **Weighted Connectivity Score** | 1 | 1 |
| **LCC Size** | 7,316,507 | 7,316,507 |
| **Number of Subgraphs** | 17,524 | 17,524 |

Both RTX-KG2 (version 2.10) and ROBOKOP KG (versioned *via* git hash `30fd1bfc18cd5ccb`) required little preprocessing as both are distributed in KGX-format and leverage Biolink Model as their data model. As ROBOKOP KG contained boolean attributes describing ontological root for diseases (e.g. MONDO_SUPERCLASS_CANCER) or drugs (e.g. CHEBI_ROLE_METABOLITE), ontologically related attributes were collapsed into a single categorical superclass attribute to reduce computational requirements for processing ROBOKOP KG. Conversely, PrimeKG (version 2.1) required significant preprocessing because it is not distributed in a KGX compatible format and does not adhere to the Biolink Model framework. To preprocess it, we first joined node indices with drug and disease feature tables to extract external identifiers which were then converted into CURIEs through prefix concatenation (e.g. X identifier from NCBI-> NCBIGene:X). Those CURIEs were then used as primary identifiers for node and edge files. As the existing PrimeKG groupings of clinically similar Mondo terms into single entities violate the Biolink Model, grouped diseases with MONDO_grouped label were split into multiple Mondo local IDs and then concatenated with the appropriate “MONDO:” prefix. Lastly, we manually mapped PrimeKG node and edge types to the Biolink model. We determined the appropriate Biolink predicate by evaluating Biolink heuristics together with the original edge type and associated attributes characterizing the relationship.

#### Ingestion

To ensure that all upstream networks are compatible with the integration pipeline and ready for downstream processing, each preprocessed KG was then validated in the ingestion pipeline (Fig. 1a). During ingestion, each upstream KG is verified for the presence of attributes relevant to downstream processing such as *id* or *category*. Additionally, the format of primary identifiers for nodes and edges is checked to match CURIE format across all upstream KGs.

#### Integration

The integration pipeline involves three distinct stages of processing the validated KGs: schema transformation, normalization of CURIE identifiers, and unification of KGs into a single network.

##### Schema Transformation

Biomedical KGs are constructed by ingesting disparate databases for different use cases, leading to systematic variations in data schemas. To achieve interoperability on the schema-level, each upstream KG underwent schema standardization to minimize data redundancy and maximize consistency across properties and data types. Attributes shared across KGs were transformed and mapped to EC-KG schema. Attributes in the upstream KGs which were not part of EC-KG schema were dropped due to their source pipeline specificity and data redundancy: for instance Mondo and CHEBI ontology superclasses attributes in ROBOKOP KG were dropped as they are represented by their respective ontologies and CURIEs. Once harmonized, we created an *upstream_data_source* attribute to track upstream network’s provenance, creating a complete EC-KG schema for nodes and edges (Tables 4,5).

##### Normalization

As biomedical networks can represent the same concepts using different ontologies and identifiers, all entities need to be resolved to a set of canonically accepted identifiers to prevent network fragmentation (Fig. 1b). For this purpose, we leveraged the Babel Node Normalizer API,^32^ which exposes identifier cliques created by the Babel pipeline, systematically processes equivalent information from a wide array of biomedical databases, and selects a preferred identifier following a source preference identifier by the Biolink Model.^32^ The Babel API was utilizing conflation, a controlled mechanism for grouping concepts closely related to each other. This conflation was applied to structurally similar chemical compounds, such as different salts or doses of the same drug compounds, and protein-encoding genes with the proteins they encode, simplifying harmonization across databases. In cases where normalization failed or a canonical CURIE was already in use, the original identifier was kept.

During node normalization, the API does not return the most specific category in Biolink Model hierarchy. To ensure nodes’ category attributes accurately represent the entity post-normalization, the most specific category in the Biolink hierarchy was retrieved from *all_categories* attribute. For instance, given *Biolink:ChemicalEntity* and *Biolink:NamedThing, Biolink:ChemicalEntity* was selected. Each preprocessed KG was normalized individually using Babel (v2.4.1) to preserve source level-integrity. Provenance was maintained by generating a new attribute with the original suffix. Successfully normalized nodes (i.e. entities for which API returned non-Null CURIe) are flagged with a *normalization_success* indicator to enable post-hoc computation of normalization rate pre-aggregation on a source-level.

##### Network Unification

To enable discovery of novel pathways across upstream networks, normalized KGs were integrated into a single unified component (Fig. 1c). The unification step was designed to preserve source specific information while minimizing redundancy in attribute fields. Node merging was conducted based on the ID field. To ensure provenance on the network- and source-level, we performed edge merging using the subject, predicate, and object fields as well as the upstream data source and primary knowledge source as composite identifiers.

When duplicate nodes or edges were identified, source-specific attributes were aggregated to retain provenance across KGs. Attributes intrinsic to the identifier, such as description for nodes or relationships direction qualifiers were then de-duplicated. For instance, when a node corresponding to aspirin is present in both PrimeKG and ROBOKOP KG, the node gets merged into a single entity with expanded *upstream_data_source* attribute tracking both PrimeKG and ROBOKOP KG as upstream KGs. However, attribute *all_categories* is only retained from one upstream KG as it is an identifier specific attribute extracted during normalization.

Biomedical KGs occasionally contain floating, standalone nodes resulting from indexing requirements or ontology design based on open-world assumption.^53^ For the purpose of drug repurposing, we consider nodes with no edges as entities with no biomedical knowledge attached to them and therefore drop them as the last step of network unification. This minimizes data network fragmentation and produces EC-KG, a unified network of normalized biomedical knowledge graphs (Fig. 1d).

##### Network Quality Metrics

To characterize the structural integrity, ontological grounding, and normalization fidelity of EC-KG, we computed a suite of metrics spanning four levels of aggregation: per-edge, per-node, per-source, and graph-global. Throughout, we denote the unified graph as *G* = (*N*, *E*), where *N* is the set of nodes and *E* the set of directed, predicate-typed edges. The set of drugs and diseases targeted for drug repurposing predictions (see Core Drugs and Diseases for Repurposing) is denoted *C* ⊆ *N*, with *N_EC_* = |*C*|, and is referred to below as ‘EC entities’

##### Ontological classification

Every edge in *E* was classified as either an instance-level (ABox) or concept-level (TBox) assertion using the Biolink Model predicate hierarchy. Specifically, an edge with predicate *p* was assigned to ABox if *p* descends from *Biolink:related_to_at_instance_level*, and to TBox if *p* descends from *Biolink:related_to_at_concept_level*. ABox and TBox edge counts were tabulated per primary knowledge source to summarize the relative contribution of factual versus terminological structure across upstream resources.

At the node level, we additionally compute an ontological connectivity indicator to distinguish nodes participating in at least one terminological assertion from bare instance nodes lacking schema-level context. For each node *n* ∈ *N*, is_ontology_connected(*n*) = 1 if ∃ *e* ∈ *E_TBox_* such that *n* ∈ endpoints(*e*); 0 otherwise.

##### Connectivity structure

Connected components of the undirected projection of *G* were enumerated using the GraphFrames library^54^. The output was reconciled by re-ranking components in descending order of size such that the largest connected component (LCC) always carries component_id = 0. Let *S* = {*S_1_*, *S_2_*, …, *S_k_*} denote the set of components, with *S_LCC_* = arg max*_Sᵢ ∈ S_* |*S_i_*| and *N_LCC_* = |*S_LCC_*|.

For each component *S_i_*, we record its size |*S_i_*|, an MD5 hash computed over the lexicographically sorted node IDs in *S_i_*, and the per-component counts of core drugs and diseases, and remaining nodes:

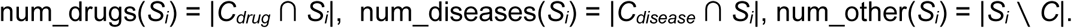

The total number of components |*S*| is retained as a graph-global summary statistic.

##### EC Core Entity connectivity

To quantify how well the curated drug and disease lists are embedded in the unified graph, we computed two summary statistics, separately for drugs only, diseases only, and the combined set. Let *C_i_* = *C* ∩ *S_i_* denote the EC core nodes in component *i*, and *C_LCC_* = *C* ∩ *S_LCC_* those in the largest component. Both metrics are treated as the graph-global summaries for KG quality assessment.

###### LCC fraction

The LCC fraction measures the share of curated drugs and diseases residing in the main connected mass of the graph:

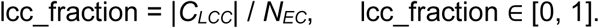

A value approaching 1 indicates that nearly all curated entities are mutually reachable, whereas lower values reflect network fragmentation across disconnected sub-structures.

###### Weighted connectivity score

The weighted connectivity score refines the LCC fraction by additionally penalizing drugs and diseases that, while present, reside in small isolated components:

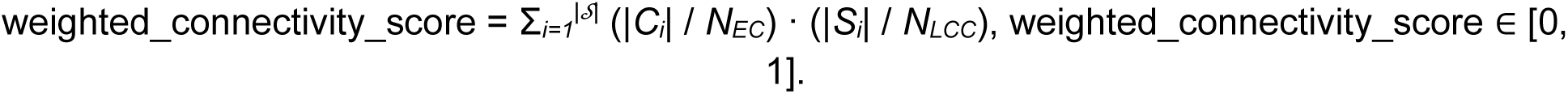

The first factor weighs each component by its share of EC entities; the second weighs each component by its size relative to the LCC. The score approaches its upper bound when all drugs and diseases lie within the LCC and approaches zero as they are dispersed into progressively smaller components.

### Infrastructure

Integration of heterogeneous biomedical knowledge graphs requires significant memory resources and scalable computational infrastructure. To address this, we used Apache PySpark (v3.5) for distributed computing within the Kedro framework and Argo Workflows (v3.7.0) for orchestration on a Kubernetes cluster. The parallel pipeline execution utilized virtual machines with RAM memory between 32-128 GB and GCS storage for data layering architecture, with a run-time of approximately 100 minutes. For scientific reproducibility, EC-KG construction can also be executed sequentially on a single virtual machine with a minimum of 128 GB of RAM and 30 GB of disk space, with the run-time of approximately 200 minutes. Both infrastructure and compute layers required for each pipeline were codified with Terraform.

### FAIR design of EC-KG

To maximize the reuse of EC-KG by researchers and computational agents, its integration and release pipelines were guided by FAIR (Findable, Accessible, Interoperable, Reusable) data principles. EC-KG releases are automatically published to the Hugging Face (HF) dataset repository with associated metadata and DOI, with data schema described following Croissant formatting^55^ for findability. To ensure accessibility, EC-KG is distributed through HF Datasets and Dask Python libraries, enabling programmatic access through HF API over HTTPs protocol. Interoperability is incorporated in the integration pipeline through schema standardization, identifier normalization, and Biolink Model compatibility (see Integration section). EC-KG integration, curation and harmonization is licensed under the Creative Commons Attribution 4.0 International License for reusability, with licenses of all upstream knowledge sources still applying.

### Data Records

EC-KG is a heterogeneous, directional biomedical KG developed as a part of Every Cure’s drug repurposing platform, which begins with identifying promising drug repurposing ideas and then evaluating these ideas in the laboratory and/or clinical trials and advancing them to patient use.EC-KG has already been successfully used in drug repurposing research with the RTX-KG2-ROBOKOP KG subgraph being used to develop an ML classifier identifying Botulinum Toxin A as a candidate for Major Depressive Disorder, as well as to validate repurposing of Lenalidomide and Dexamethasone against Refractory Rosai-Dorfman Disease.^47,48^ It contains over 7 million nodes and 81 million edges with associated attributes. The knowledge base embeds ontological richness across a diverse set of biological networks such as protein-protein interactions, drug-pathway relationships and disease-phenotype associations (Fig. 2). Its FAIR aligned design supports benchmarking of KG-based computational drug repurposing methods across integrated networks, enabling the development of scalable approaches for knowledge discovery. EC-KG is composed of over 60 unique node types and 91 unique edge types derived from the hierarchical Biolink Model for predicates and node types, allowing for queries and subgraph extraction at selective levels of biomedical granularity.

**Figure 2.**
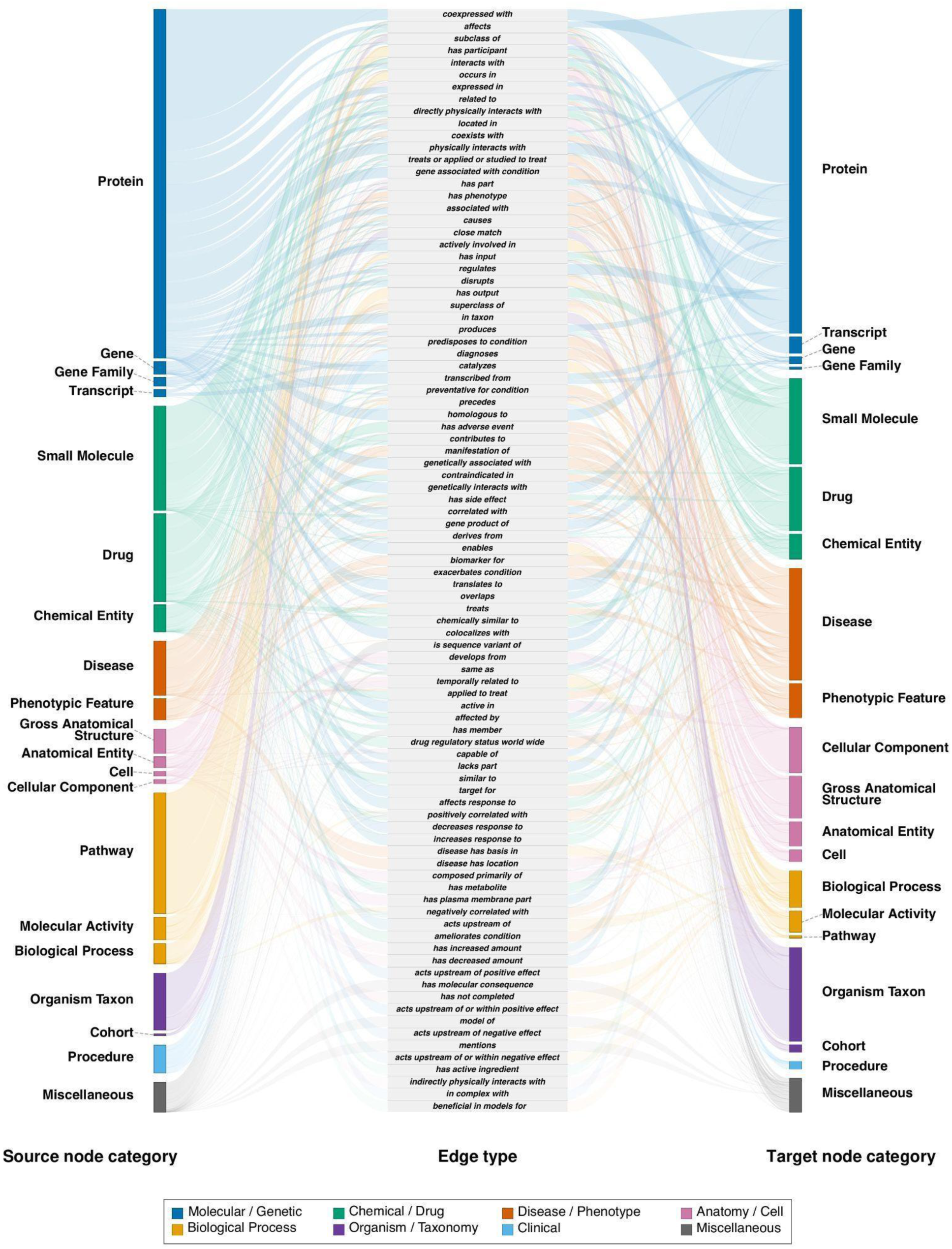
Sankey diagram of EC-KG edge flows by Biolink Model predicate. Source node category (left), edge type (middle), and target node category (right). Bars represent the quantity of nodes within each node category, colored by semantic supergroup (dark blue: Molecular/Genetic, yellow: Biological Process, green: Chemical/Drug, orange: Disease/Phenotype, pink: Anatomy/Cell, purple: Organism/Taxonomy, light blue: Clinical, gray: Miscellaneous); the central column lists Biolink Model predicates connecting them. Ribbon height is proportional to edge count for each source–predicate–target combination. Node categories that contribute less than 0.5% of total edges as both source and target are collapsed into a single Miscellaneous bar to preserve legibility.

EC-KG was created through a parallelizable integration of PrimeKG, RTX-KG2 and ROBOKOP KG into a single biomedical knowledge graph. Despite all three upstream networks being built to represent overlapping biomedical systems, each KG contributes a significant amount of unique information as only a moderate proportion of nodes are resolved into the same entity while retaining high normalization rate across the KGs (Fig. 3a,b). High normalization rate indicates that the majority of identifiers are mapped to Babel identifier cliques,^32^ and therefore information contributed by each KG is not duplicative. While RTX-KG2 is the single largest contributor of unique information, all upstream networks capture the fundamental mechanistic entities such as drugs, proteins and diseases (Fig. 3c). The integration framework is extensible and can integrate additional knowledge graphs, including large-scale resources such as SPOKE, as well as other Biolink Model compatible KGs. The pipeline also supports extraction of targeted subgraphs through heuristic definitions based on the Biolink Model and KG level attributes.

**Figure 3.**
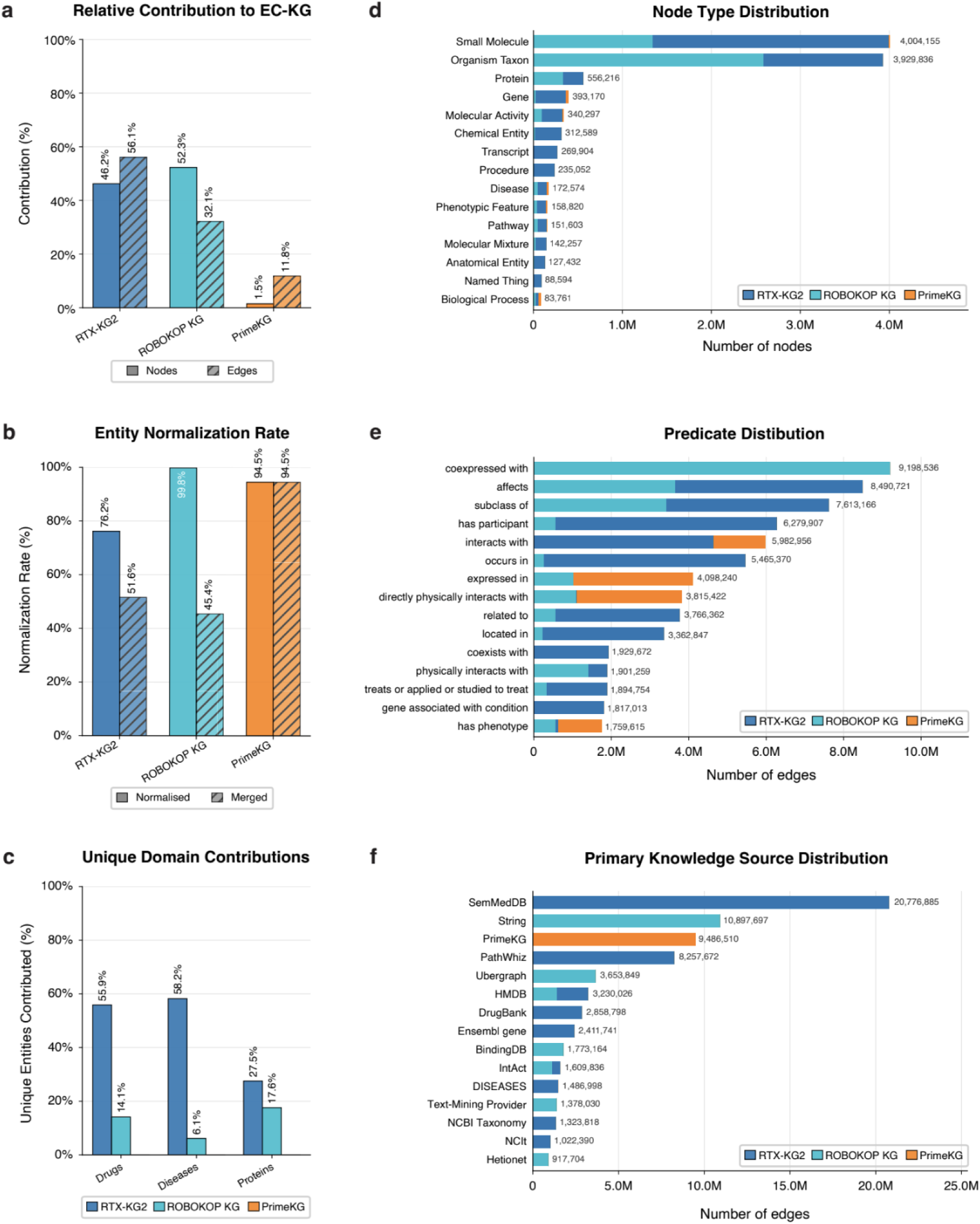
Integration statistics and distributional characteristics of the three upstream knowledge graphs comprising EC-KG. **(a)** Total share of nodes and edges contributed by each upstream KG (RTX-KG2, ROBOKOP KG, PrimeKG) relative to the unified network; PrimeKG’s disproportionately high edge share relative to its node share reflects its original Mondo grouping strategy, discussed in Methods. **(b)** Normalization and merge rates per upstream KG, showing the percentage of nodes successfully resolved to a Babel identifier clique (Normalised) versus the percentage subsequently merged with nodes from another upstream KG during unification (Merged). **(c)** Contribution ratio of each upstream KG to unique drug, disease, and protein nodes in EC-KG, calculated as the fraction of entities of each type contributed exclusively by that KG. **(d)** Distribution of the top node types in EC-KG by count, stacked and colored by upstream source. **(e)** Distribution of the top Biolink predicates in EC-KG by edge count, stacked and colored by upstream source. **(f)** Distribution of the top primary knowledge sources in EC-KG by edge count, stacked and colored by upstream source. Together, panels D–F illustrate that while foundational entity and predicate types are broadly shared across upstream networks, each KG contributes a distinct profile of primary knowledge sources, underscoring the complementarity of the three constituent graphs.

EC-KG schema is described in Table 3 and 4 for nodes and edges, respectively. The attributes include features for provenance tracking and knowledge mining approaches, as well as specific drug repurposing applications, such as qualifiers describing the nature of relationships or experimental parameters for binding affinity. While the majority of the attributes are derived from the Biolink Model, EC-KG expands the schema with features such as *upstream_data_source* for multi-level provenance, or *num_references* to characterize the number of claims supporting the edge. The use of standardized CURIEs as primary identifiers together with Biolink-derived attributes such as category and predicate forms the standardization layer of EC-KG and ensures compliance with other KGX-compliant networks.

**Table 3.** Node attributes in EC-KG. Attributes support cross-database interoperability, ontological classification, and provenance tracking.

| Attribute | Description | Example |
| --- | --- | --- |
| **Id** | CURIE identifiers represent biomedical entities. | *NCBIGene:92482* |
| **Name** | Canonical name associated with the CURIE identifier, provided by Babel. | *BBIP1* |
| **Description** | Description of the biomedical entity, provided by upstream knowledge source. | *The BBSome complex is thought to function as a coat complex required for sorting of specific membrane proteins to the primary...* |
| **Category** | Biolink Category associated with the CURIE, provided by Babel | *Biolink:Protein* |
| **All Categories** | All Biolink ancestor categories associated with the Id identifier. | [*"Biolink:Gene", "Biolink:GeneOrGeneProduct", "Biolink:PhysicalEssence", "Biolink:OntologyClass", , "Biolink:Protein"...]* |
| **Equivalent Identifiers** | Identifiers equivalent to id. | [*"UniProtKB:A8MTZ0", "ENSEMBL:ENSG00000214413", "HGNC:28093", "NCBIGene:92482", ...]* |
| **International Resource Identifiers** | URL for identifier CURIE resolution. | *https://identifiers.org/ncbigene:92482* |
| **Upstream Data Source** | List tracking provenance of upstream networks used for integration. | [*"primekg", "rtxkg2", "robokop"]* |
| **Publications** | DOI and PMID identifiers representing publications where identifier was mentioned | [*"DOI:10.1136/jmedgenet-2013-101785", "PMID:15164054", "DOI:10.1101/gr.2596504" ...]* |

**Table 4.**
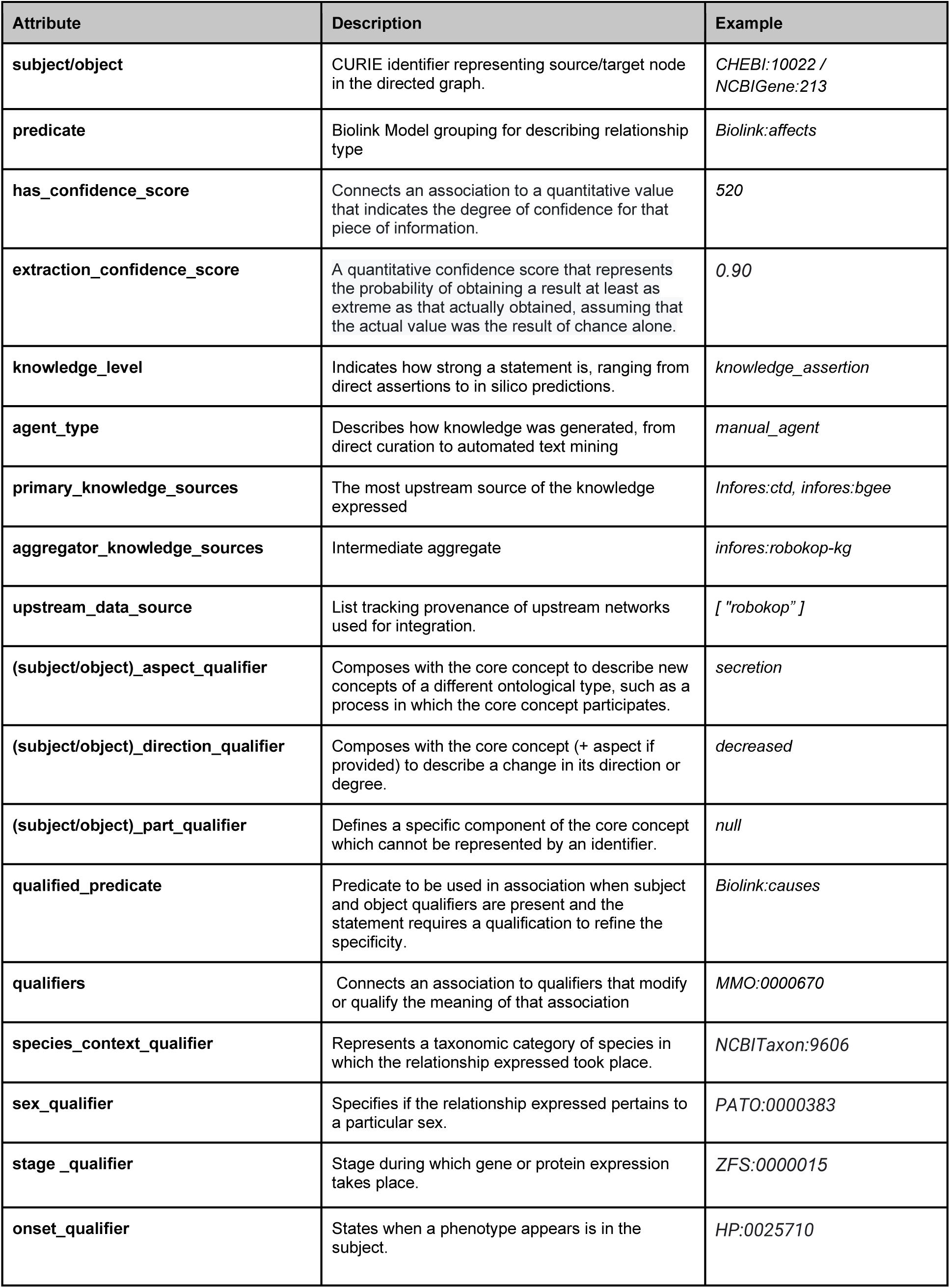
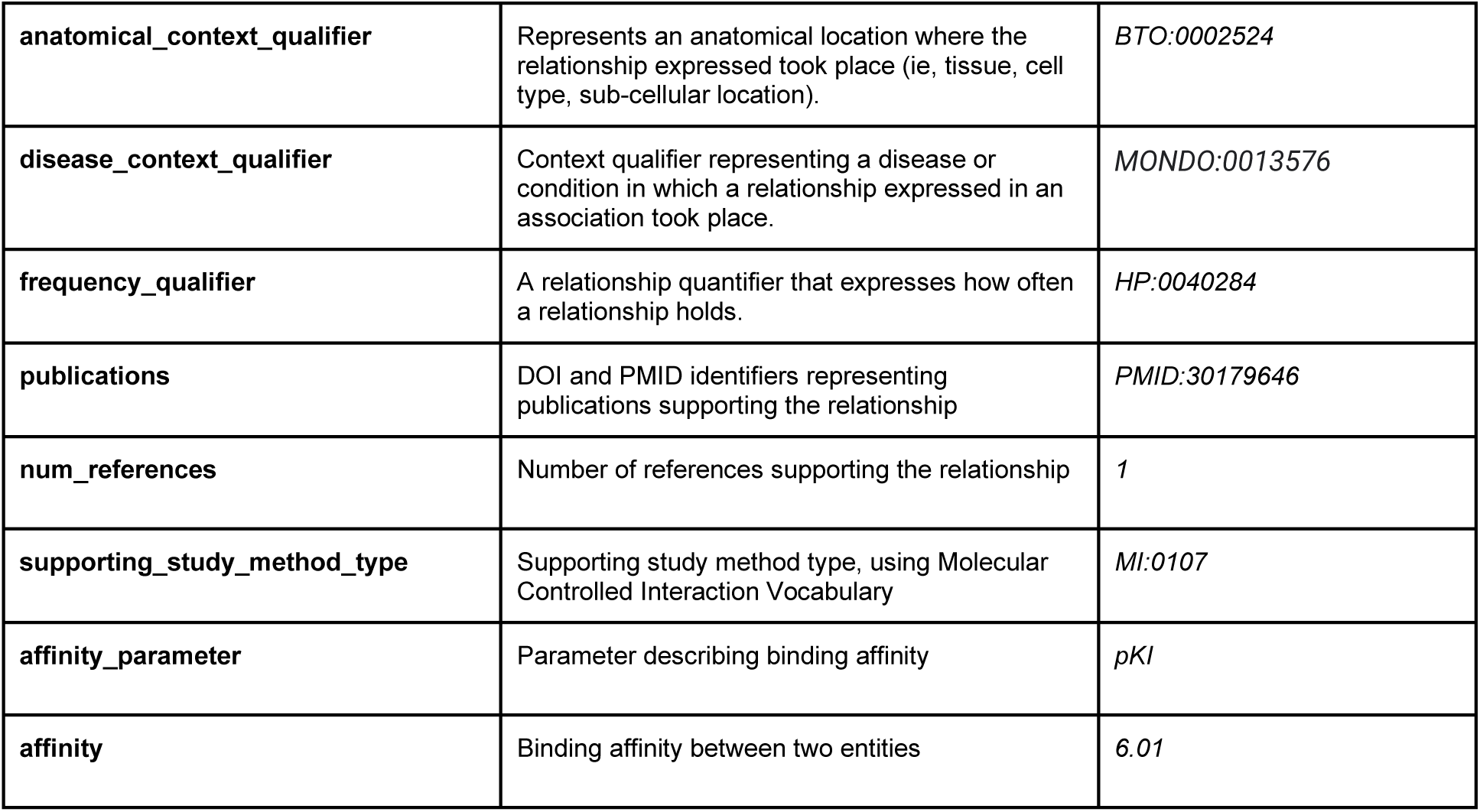
Edge attributes in EC-KG. Because no single subject-predicate-object triplet contains all attributes simultaneously, the examples are compiled from different edges across the network to illustrate the full range of available attributes in EC-KG.

EC-KG was designed with FAIR principles as explicit design requirements, with its schema, standardization strategy and attributes mapping to each FAIR subprinciple (Table 5). It is findable through the HF dataset registry where it is indexed with persistent identifiers and metadata, and accessible programmatically over HTTPs via the HF API. Interoperability is achieved through Biolink Model compatibility, Babel identifier normalization, and provenance-oriented edge attributes. Finally, adherence to well-established Biolink Model and KGX formatting, and clear licensing ensure that EC-KG meets established community standards for KG sharing and reuse. Together these features increase the transparency and trustworthiness of EC-KG for drug repurposing research.

**Table 5.**
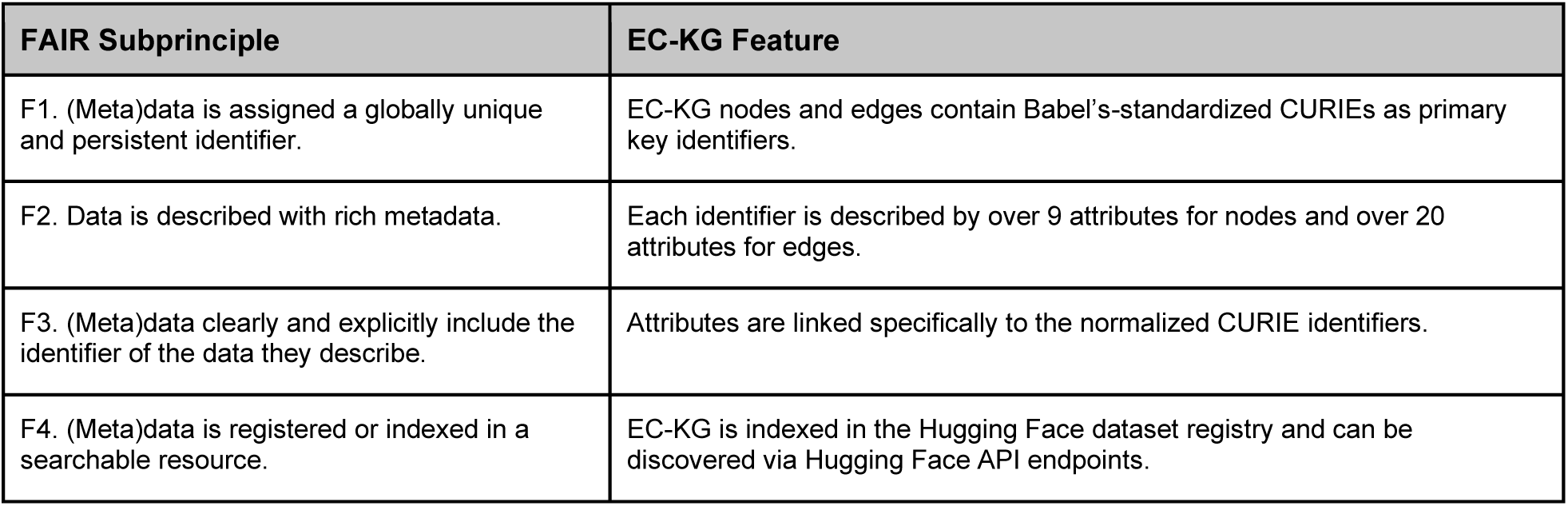
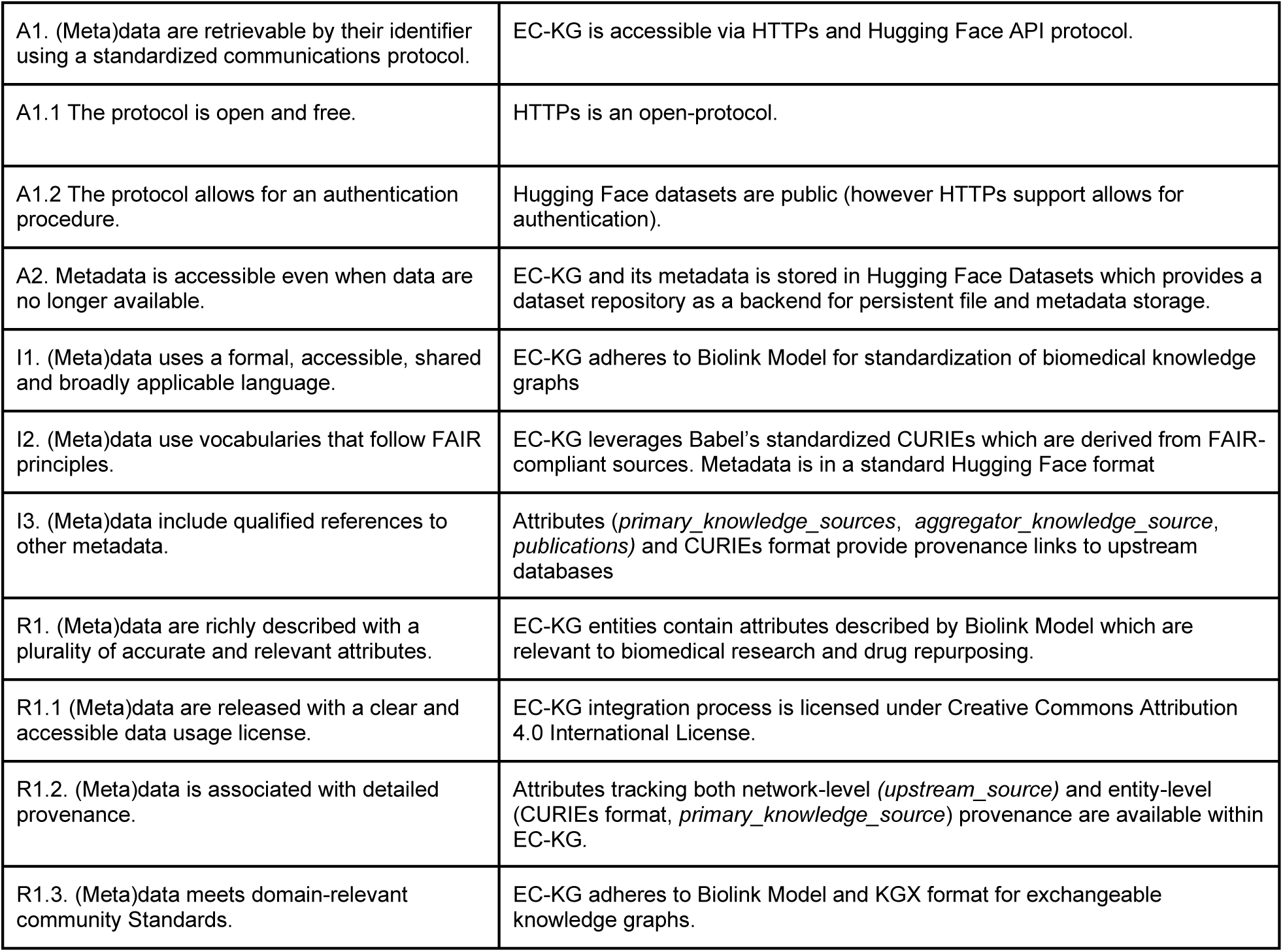
FAIR subprinciples^37^ addressed by EC-KG design. Through multi-level provenance tracking, integration of interoperability tools like Biolink Model or Babel, and Hugging Face hosting, EC-KG addresses all FAIR guiding principles.

### Technical Validation

In this section, we demonstrate how EC-KG’s FAIR-centric design addresses gaps in provenance, interoperability, and completeness. First, we validate multi-level provenance tracking within EC-KG through a comprehensive analysis of aggregated primary knowledge sources, quantifying source granularity, completeness, and associated diversity of biomedical ecosystems feeding into the network. Second, we demonstrate how the Biolink Model and the normalization strategy enable extraction of fully interoperable subgraphs from EC-KG for data-centric benchmarking. Third, we show how unification of different biomedical networks leads to the emergence of novel metapaths and mechanistic pathways relevant for drug repurposing. Last, we demonstrate how EC-KG can be used for computational drug repurposing by predicting drug-disease treatment relationships and evaluating them on curated off-label indication sets.

### Knowledge Source Aggregation

To validate multi-level provenance tracking capabilities of EC-KG, we analyzed knowledge sources aggregated from the upstream networks. First, we quantitatively assessed the knowledge source contributions on the network and edge-level to assess unique and overlapping contributions across the integrated network. Second, we conducted a detailed assessment of the underlying knowledge sources to evaluate the diversity of biomedical information introduced by the aggregation strategy.

In total, EC-KG integrates 95 different knowledge sources through network-level aggregation. Although PrimeKG ingests over 20 different primary sources, we treated the integrated version of the graph as a distinct knowledge source (Fig. 3f). This is due to the lack of provenance-tracking on the node and edge-level in PrimeKG, as well as additional custom modifications applied to its primary sources such as node merging of Mondo^51^ diseases or selection of only highly-expressed human genes from the Bgee database.^58^ As a result, only 15 out of 95 aggregated sources (16%) overlap between RTX-KG2 and ROBOKOP KG, with PrimeKG being a distinct entity with no formal overlap. However, if all PKS used to construct PrimeKG are considered separate, then 20 out of 94 sources overlap, as all PKS of PrimeKG are shared within ROBOKOP KG and RTX-KG2. Although a significant proportion of entities overlap across the upstream KGs, the majority of subject-predicate-object triplets are unique to a single source. This demonstrates that each graph contributes distinct new knowledge to nodes shared across all KGs.

The shared PKS include gold-standard databases such as Reactome^59^ (edges N=706,342), IntAct^60^(N=1,609,013) and HMDB^61^ (N=3,230,026), which capture fundamental drug-target-disease relationships enriched with pathway and metabolic data. This overlap forms a high-confidence backbone for EC-KG, which the distinct sources from each upstream network then extend with different layers of knowledge.

The unique contributions provided by RTX-KG2 include literature-mined associations together with curated metabolic pathways and clinical drug information through sources like SemMedDB^62^, PathWhiz^63^ or NCIT (Table 6). The ontological layer encoded within RTX-KG2 through the use of extensive upper-level ontologies (e.g. RO^64^, IAO^65^) together with core biomedical entities enables fine-grained semantic reasoning, creating the ontological skeleton of EC-KG.

**Table 6.** Top 10 Primary knowledge sources overlapping within EC-KG and unique to upstream knowledge graphs. PrimeKG is reported as a single knowledge source due to design heuristics and lack of entity-level provenance tracking.

| Unique ROBOKOP KG |  | Unique RTX-KG2 |  | Unique PrimeKG |  |
| --- | --- | --- | --- | --- | --- |
| PKS | count | PKS | count | PKS | count |
| infores:string | 10,896,709 | infores:semmeddb | 20,304,934 | infores:primekg | 8,789,126 |
| infores:ubergraph | 3,653,161 | infores:pathwhiz | 8,232,041 |  |  |
| infores:bindingdb | 1,753,285 | infores:ensemble-gene | 2,411,683 |  |  |
| infores:text-mining-provider-targeted | 1,248,746 | infores:ncbi-taxonomy | 1,323,818 |  |  |
| infores:hetionet | 917,698 | infores:ncit | 1,016,881 |  |  |
| infores:bgee | 866,863 | infores:mesh | 831,618 |  |  |
| infores:panther | 682,916 | infores:pr | 699,021 |  |  |
| infores:lincs | 544,881 | infores:umls-metathesaurus | 496,498 |  |  |
| infores:pharos | 389,056 | infores:chebi | 362,802 |  |  |
| infores:hpo-annotations | 252,571 | infores:fma-umls | 241,677 |  |  |

ROBOKOP’s unique primary knowledge sources include STRING,^66^ BindingDB,^67^ and LINCS68 (Table 6). STRING provides extensive protein-protein interactions (PPI) data, including over 9 million protein coexpressions relationships. LINCS enriches this network through transcriptomic and genetic perturbation networks, providing regulatory relationships connecting the proteins with other proteins or small molecules. Supplemented with BindingDB’s protein-ligand affinities, the integration creates an interconnected biomedical network optimized for biomedical research within EC-KG.

PrimeKG’s unique contributions include a subset of high-confidence drug-disease associations with the focus on clinical applications. Its translation to Biolink model leads to substantially richer predicate vocabulary, significantly expanding semantic expressiveness of PrimeKG while also enabling interoperability with remaining networks. This creates a curated precision medicine layer in EC-KG.

Each upstream KG contributes a distinct and complementary set of data ecosystems and metapaths to EC-KG, reflecting the complementarity between the ontological focus of RTX-KG2, the biological layer of ROBOKOP KG, and the clinical focus of PrimeKG (Fig. 3d, 3e). EC-KG has a TBox-to-ABox ratio of 0.84, indicating that the knowledge encoded in the network contains primarily instance-level relationships describing real-world biomedical relationships while preserving useful ontological structures for reasoning with many of the terminological relationships. Such a knowledge source contains a higher diversity of pathways, which can involve mechanistic pathways relevant to drug repurposing hypothesis generation. (See “Emergence of Novel Context”).

Multi-level provenance is captured at both the network and source level through rich edge metadata. For example, the interaction between Ezogabine, a positive allosteric modulator of KCNQ2 potassium channels, and the KCNQ2 protein itself is present in both RTX-KG2 and ROBOKOP, but sourced from different databases, with 2 and 4 supporting references respectively (Fig. 5). Through our network unification strategy, EC-KG successfully aggregates complementary metadata at multiple scales: over 505,117 edges were enriched with overlapping publication records while over 3,025,695 edges were supported by independent primary knowledge sources. Retaining this provenance enables downstream use cases such as for confidence score calculations, evidence-weight modelling and LLM-assisted literature review.

The integration also improves coverage of drugs and diseases essential for drug repurposing; as shown in Table 2, none of the source KGs fully covers all drugs and diseases present in the core drug and disease list, with PrimeKG having less than 80% of the drugs present within its network. Upon unification of the three networks, coverage of drugs and diseases increases, enabling drug repurposing discovery which would be limited by a single KG. Furthermore the connectivity metrics show all curated drugs and diseases are within the largest graph component, confirming a good level of integration and lack of network fragmentation of the main subgraphs. This enables computational methods to discover new pathways between drugs and diseases, which is a key inferential step in computational drug repurposing approaches such as pharmacophenomics.

The core drugs and diseases in EC-KG have rich biomedical neighborhoods enabling drug repurposing research with their emphasized drug-target-disease metapaths (Fig. 4). The disease neighborhoods integrate phenotypic networks through ontologies such as the Human Phenotype Ontology (HP)^56^ and the Disease Ontology (DOID)^57^, embedding a rich disease-target network with phenotypic associations, enabling target-driven and phenotype-driven drug repurposing approaches for both common and rare diseases which are captured by the Mondo ontology. The drug neighborhoods capture extensive coverage of drug-drug interactions provided by databases such as ChEMBL or ChEBI, making it suitable for both mono- and polypharmacy modelling.

**Figure 4.**
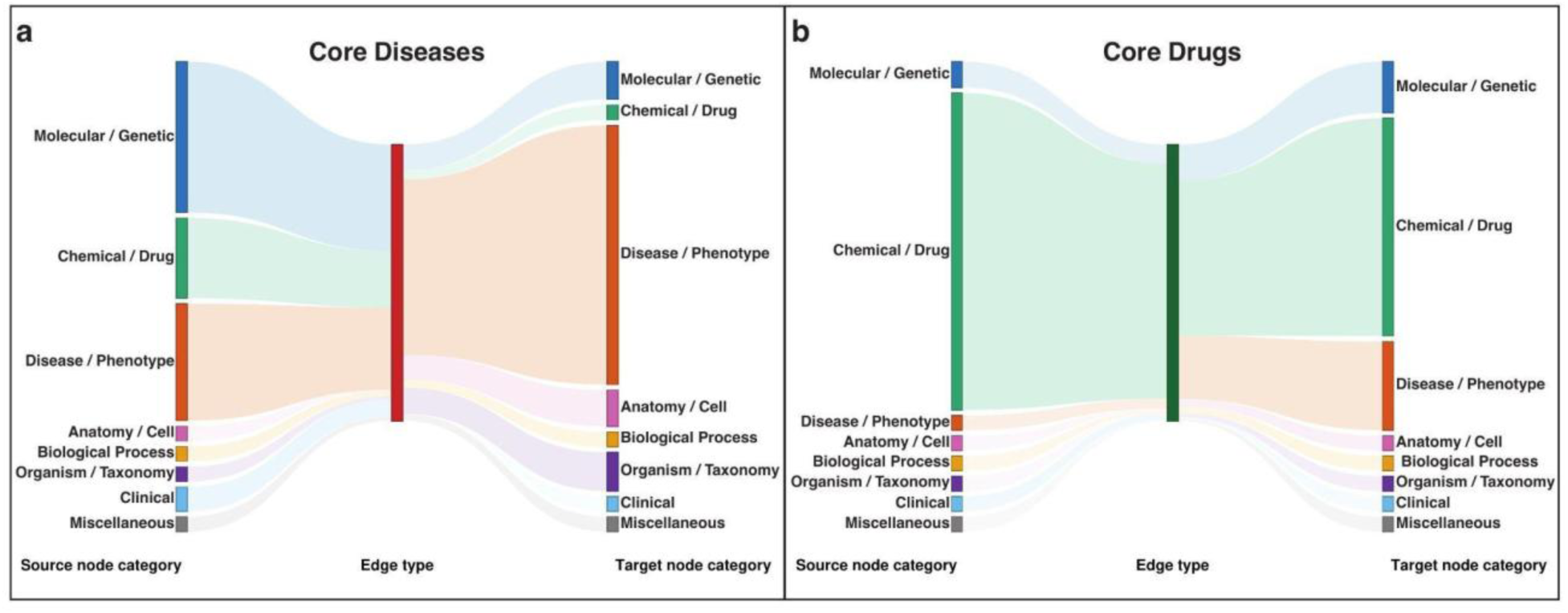
Category-level neighborhood composition of core drug and disease entities in EC-KG. **(a)** Sankey diagram of the one-hop neighborhood of core disease entities, showing source node categories (left) flowing through an aggregate “Core Disease” node (center, red) to target node categories (right). **(b)** Equivalent diagram for core drug entities (center, dark green). In both panels, source and target bars represent Biolink node categories, colored by semantic supergroup (blue: Molecular/Genetic, green: Chemical/Drug, orange: Disease/Phenotype, pink: Anatomy/Cell, yellow: Biological Process, purple: Organism/Taxonomy, light blue: Clinical, gray: Miscellaneous), with ribbon width proportional to the number of edges connecting each category to the core entity set.

**Figure 5.**
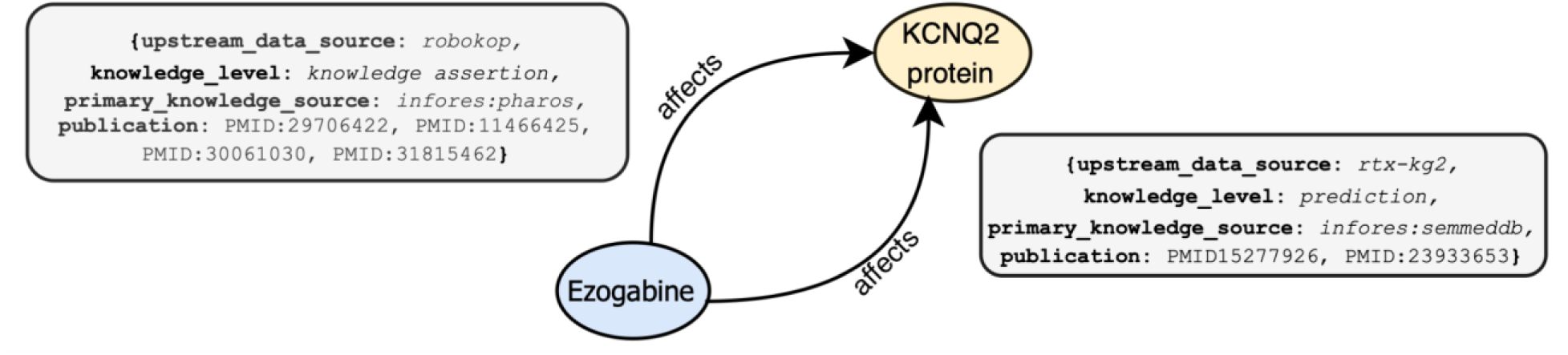
Attribute complementary for Ezogabine (CHEBI:68584) and KCNQ2 (NCBIGene:3785) across different primary knowledge sources for RTX-KG2 and ROBOKOP KG.

### Interoperable Subgraph Extraction

A feature of EC-KG being harmonized to a single standardized schema and data model is the fact that EC-KG subgraphs inherit its interoperability, enabling benchmarking and method development. To demonstrate the functional interoperability, we extracted individual task-specific subgraphs underlying well-established drug repurposing or drug discovery algorithms, and subgraphs combining multiple upstream KGs (Table 7). This extraction was performed with a filtering module within the EC-KG construction framework which uses EC-KG attributes and hierarchical structure of the biolink model to define subgraphs through inclusionary and exclusionary patterns at both the node and edge level.

**Table 7.**
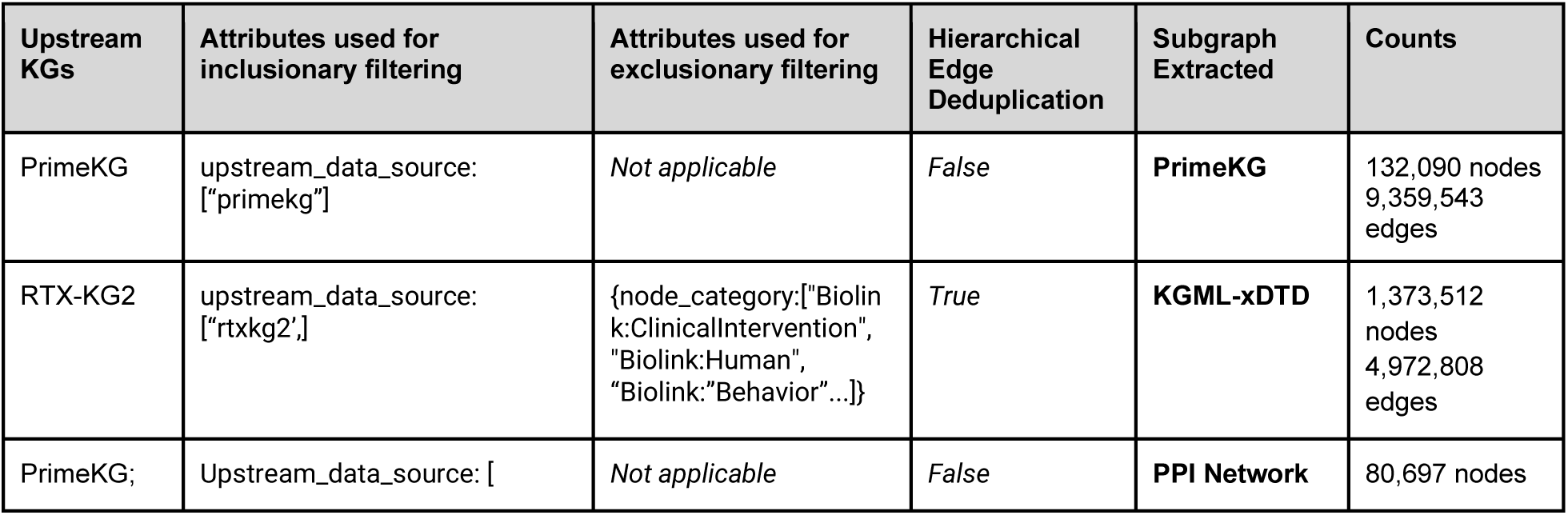
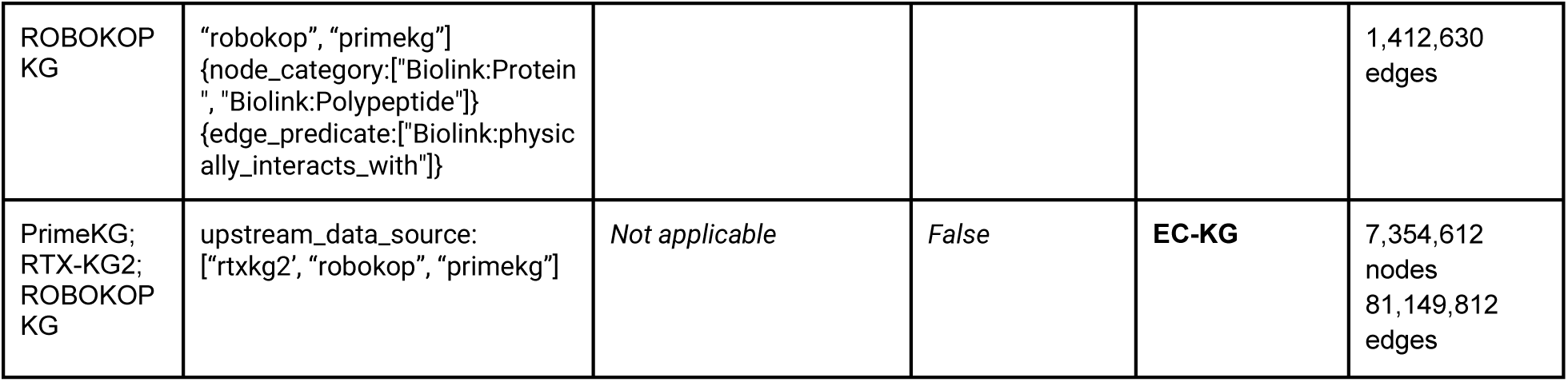
Subgraphs extracted from EC-KG using the filtering module. The subgraph extraction pipeline demonstrates EC-KGs suitability as a benchmarking graph for general drug discovery applications.

| Upstream KGs | Attributes used for inclusionary filtering | Attributes used for exclusionary filtering | Hierarchical Edge Deduplication | Subgraph Extracted | Counts |
| --- | --- | --- | --- | --- | --- |
| PrimeKG | upstream_data_source: ["primekg"] | *Not applicable* | *False* | **PrimeKG** | 132,090 nodes<br>9,359,543 edges |
| RTX-KG2 | upstream_data_source: ["rtxkg2"] | {node_category: ["Biolink: ClinicalIntervention", "Biolink: Human", "Biolink: Behavior" ...]} | *True* | **KGML-xDTD** | 1,373,512 nodes<br>4,972,808 edges |
| PrimeKG; | Upstream_data_source: [ | *Not applicable* | *False* | **PPI Network** | 80,697 nodes |
| ROBOKOP KG | "robokop", "primekg"<br>{node_category:["Biolink:Protein", "Biolink:Polypeptide"]}<br>{edge_predicate:["Biolink:physically_interacts_with"]} |  |  |  | 1,412,630 edges |
| PrimeKG;<br>RTX-KG2;<br>ROBOKOP KG | upstream_data_source:<br>["rtxkg2", "robokop", "primekg"] | *Not applicable* | *False* | **EC-KG** | 7,354,612 nodes<br>81,149,812 edges |

The first such subgraph is PrimeKG which was used for development of methods like NetMedGPT^18^ or TxGNN^17^. The original PrimeKG was constructed to have 30 distinct edge types where the majority of the edge types are represented by source and target nodes being connected by a relationship. The translation of PrimeKG into a Biolink-compatible KG expands the data model to over 87 unique triplet types. This expansion enables granular differentiation between entities, allowing for benchmarking of development of robust GNNs which frequently struggle with highly heterogenous graph topologies.

Another example is the KGML-xDTD^21^ subgraph. KGML-xDTD is a ML pipeline for drug repurposing generated by filtering RTX-KG2. Such a subgraph can be extracted by excluding entities irrelevant to mechanism-of-action predictions. As upstream KGs allow multiple edges of the same Biolink branch to connect two nodes, the filtering pipeline also allows for hierarchical deduplication, making a credible reflection of RTX-KG2 and allowing for use of EC-KG for benchmarking.

Similarly, isolated protein-protein subgraphs can be used as features to enrich the predictive power of ML models or for recommender system generation.^68^ The extraction module enables users to extract only protein-protein interactions. Search can also be limited to specific primary knowledge sources, such as STRING,^66^ or specific qualifiers which describe the nature of relationships.

The FAIR-aligned design of EC-KG together with the filtering module allows for the easy extraction of complete topologies of task-specific subgraphs as well as ROBOKOP KG, RTX-KG2, and PrimeKG in a fully interoperable format. This demonstrates EC-KG’s capability for interoperability and its utility for ML method development and comparative benchmarking (see Prediction of Drug Repurposing Candidates).

### Emergence of Novel Context

As direct drug-disease edges do not provide mechanistic context relevant to the pair, surrounding neighborhoods need to be explored to assess the contextual and mechanistic richness of drug-disease pairs encoded within the KG. Suboptimal Paths (SOPs) explore such context by extracting paths between two nodes excluding shortest one-hop connections, therefore including alternative routes between the drug and disease of interest through intermediate entities such as genes, proteins or biological processes. Such SOPs are particularly relevant for drug repurposing: if a drug and disease are not directly connected but share biologically meaningful intermediate nodes or processes, these relationships may suggest previously unrecognized therapeutic hypotheses as biologically meaningful pathways. Such pathways however might not always be visible within a single non-interoperable KG: by integrating three upstream KGs, the context is aggregated across multiple databases and ontologies, increasing the diversity of suboptimal paths and linking previously disconnected nodes. This integration increases the range of possible metapaths connecting two nodes of interest, including pathways involving biological mechanisms and evidence types represented in the graphs as well as biological explanations surrounding biologically meaningful pathways.

To demonstrate how unification of different biomedical networks leads to emergence of novel mechanistic paths, we conducted SOP analysis using the curated off-label indications. We selected a subset of pairs present across PrimeKG, ROBOKOP KG, RTX-KG2 and EC-KG and randomly sampled 100 pairs without replacement. We enumerated all simple paths up to 2 and 3 hops between each drug-disease pair using a modified depth-first search algorithm.^69^

As the unification of networks inevitably leads to an exponential increase in paths between two nodes of interest, including paths representing topological noise, we validated the quality of SOP in two ways. First, to ensure unification of the networks into EC-KG leads to genuine semantic integration rather than artificial edge densification, we computed the 2-hop SOPs for a sample of one hundred randomly rewired KGs. Then, to ensure SOPs include biomedically relevant pathways as a result of unification, we systematically examined the metapaths encoded within the SOPs through intermediate node type distribution analysis, validating the emergence of novel mechanistic paths within the new SOPs.

EC-KG provides significantly more pathways connecting drugs and diseases than constituent upstream networks, indicating a richer topological connectivity than either the standalone upstream KGs or randomly-rewired null models (Table 8). Evaluation of individual off-label indications shows that while none of the upstream KGs has consistently superior representation of all off-label pairs, their unification into EC-KG yielded the most information-rich subgraphs for the drug repurposing cases (Table 9).

**Table 8.**
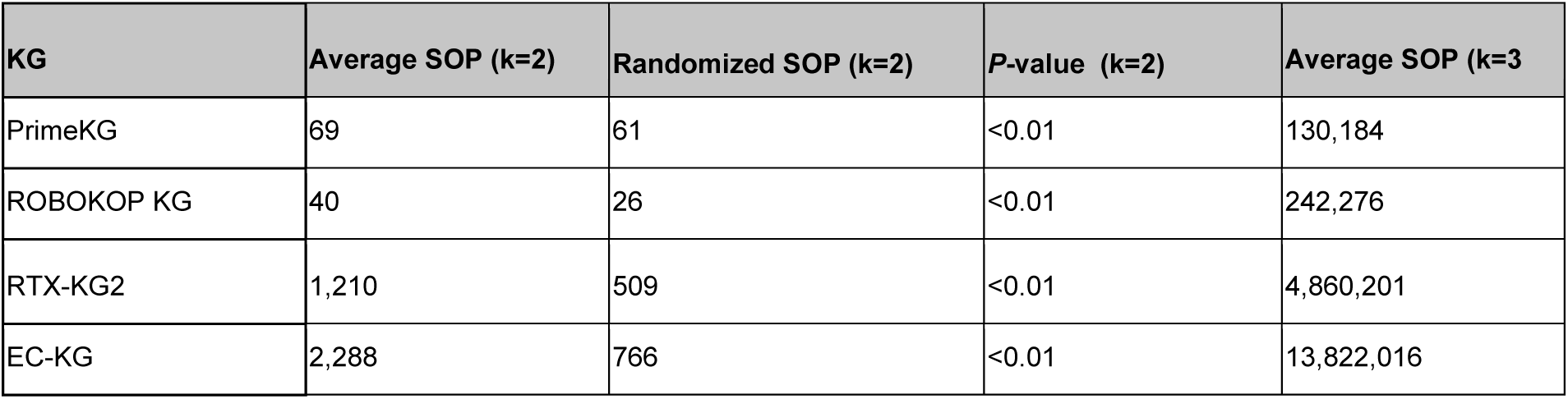
Average number of 2- and 3-hop long suboptimal paths for 100 off-label pairs present across all KGs.

| KG | Average SOP (k=2) | Randomized SOP (k=2) | P-value (k=2) | Average SOP (k=3) |
| --- | --- | --- | --- | --- |
| PrimeKG | 69 | 61 | <0.01 | 130,184 |
| ROBOKOP KG | 40 | 26 | <0.01 | 242,276 |
| RTX-KG2 | 1,210 | 509 | <0.01 | 4,860,201 |
| EC-KG | 2,288 | 766 | <0.01 | 13,822,016 |

**Table 9.** Examples of off-label indications with their corresponding SOPs across biomedical networks. The variance across all three upstream KGs shows KG complementarity and the need for KG integration for emergence of novel context.

| Drug | Disease | PrimeKG SOP (k=2) | ROBOKOP KG SOP (k=2) | RTX-KG2 SOP (k=2) | EC-KG SOP (k=2) | PrimeKG KG SOP (k=3) | ROBOKOP KG SOP (k=3) | RTX-KG2 SOP (k=3) | EC-KG SOP (k=3) |
| --- | --- | --- | --- | --- | --- | --- | --- | --- | --- |
| Bortezomib | Primary CTCL | 128 | 31 | 6 | 600 | 230,344 | 108,874 | 5,827 | 2,102,039 |
| Gemcitabine | Ewing sarcoma | 42 | 23 | 1,802 | 27,168 | 75,882 | 329,087 | 4,284,474 | 10,310,425 |
| Everolimus | Waldenstrom macroglobulinemia | 56 | 58 | 761 | 1,352 | 39,872 | 105,310 | 417,599 | 5,257,637 |
| Cladribine | T-cell large granular lymphocyte leukemia | 0 | 17 | 192 | 437 | 48 | 31,595 | 318,553 | 979,038 |
| Fondaparinux | acute coronary syndrome | 90 | 28 | 1,446 | 3,680 | 164,714 | 241,770 | 1,220,804 | 15,727,199 |

Unification of networks also leads to an increase in biomedically relevant connections between two nodes of interest. To quantify the scope of novel mechanical context emergence among the biomedically irrelevant pathways, we systematically identified metapaths in EC-KG that do not exist in any individual source KG. A path is classified as “EC-KG-unique” if at least one intermediate node or edge on the path originates from a different source KG than the endpoints’ primary source. The paths were then classified as biomedically irrelevant if they contained a node type irrelevant to mechanistic pathways (e.g. *biolink:Human*, *biolink:Food)*.

This led to identification of 348 unique paths across the 3-hop SOPs, with 45% of metapaths being considered as biomedically meaningful (Fig. 6a). The SOP traversal through EC-KG is also directed through significantly more mechanistically relevant entities such as proteins or biological pathways, demonstrating that the integration of individual KGs increases the range of biological mechanisms and evidence types represented in the graphs (Fig. b). Although the most prevalent metapaths are not mechanistically relevant, emergence of such novel contexts can be important for use cases such as clinical trial success or adverse reaction prediction.^70^

**Figure 6.**
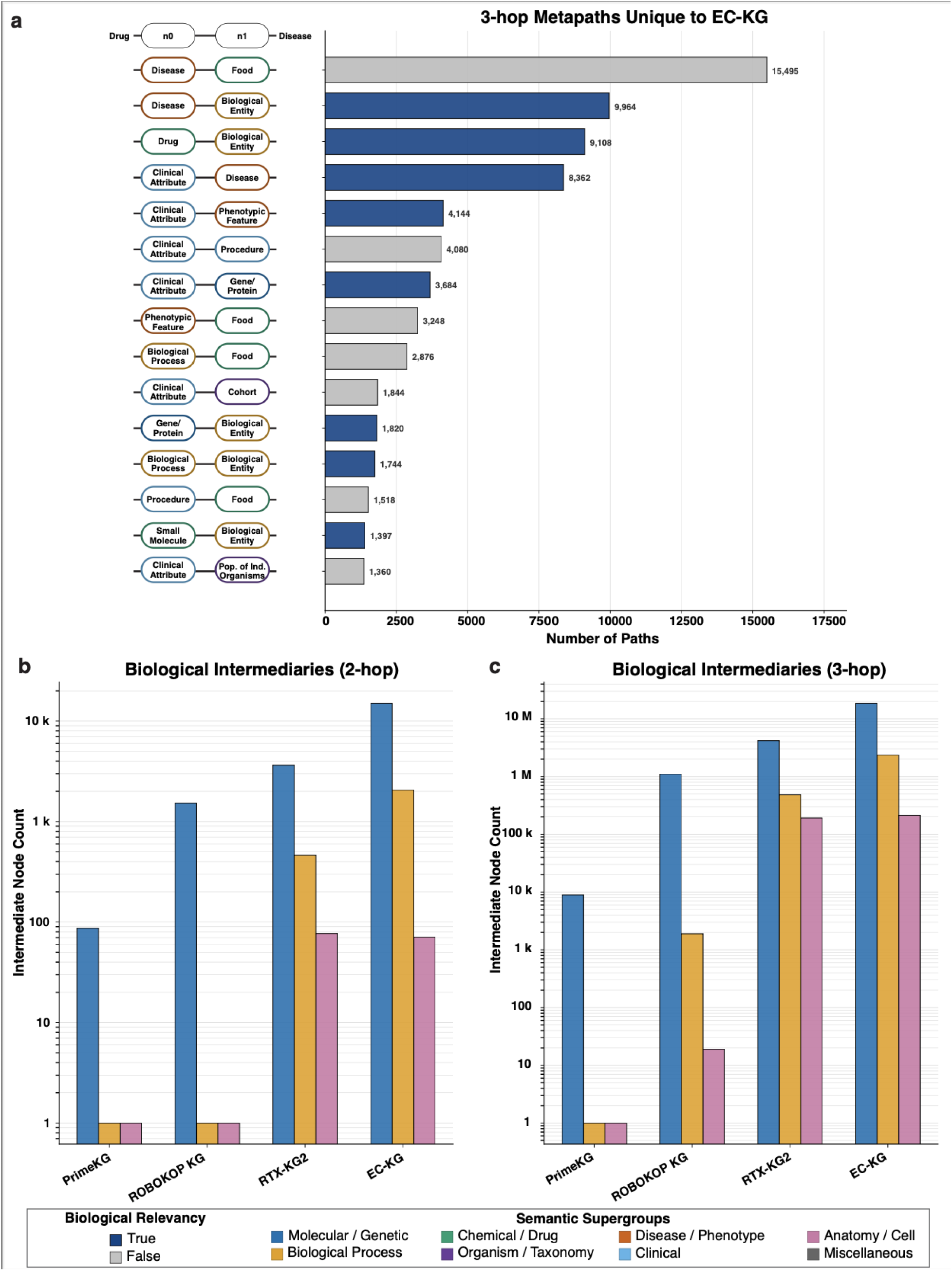
Analysis of metapaths and intermediate nodes in the generated SOPs. a) Distribution of top 15 metapaths unique to EC-KG and connecting off-label drug-disease pairs within their 3-hop SOPs. The y-axis labels display the intermediate node transitions mapping the alternative routes connecting drug-disease pair, with drug and disease nodes being omitted for readability (as they are static across all SOPs). The biomedical relevance tab distinguishes mechanistic pathways from topological artifacts. b) Distribution of key biomedical entities as intermediate nodes within 2-hop and 3-hop SOPs between repurposed drugs and diseases, showing increase in biomedically relevant entities within SOPs within EC-KG

One specific example of novel context emergence can be a SOP of Bortezomib and Primary cutaneous T-Cell Lymphoma (CTCL) (Fig. 7). Bortezomib is primarily approved for multiple myeloma patients. However, it has been clinically studied for cutaneous T-cell lymphoma and is often prescribed off-label.^71^ While a direct connection between Bortezomib and CTCL are present across all upstream KGs, many mechanistically relevant connections are missing. EC-KG provides mechanistic context emerging from integration of all upstream networks: Bortezomib is interacting with CDKN2A protein, a cyclin-dependent kinase inhibitor crucial to prevent transformation of T-cells in CTCL.^72,73^ Both of these mechanistic targets emerge only upon integration of upstream KGs, indicating complementary evidence streams which may identify novel therapeutic connections which would normally be missed by querying a single network.

**Figure 7.**
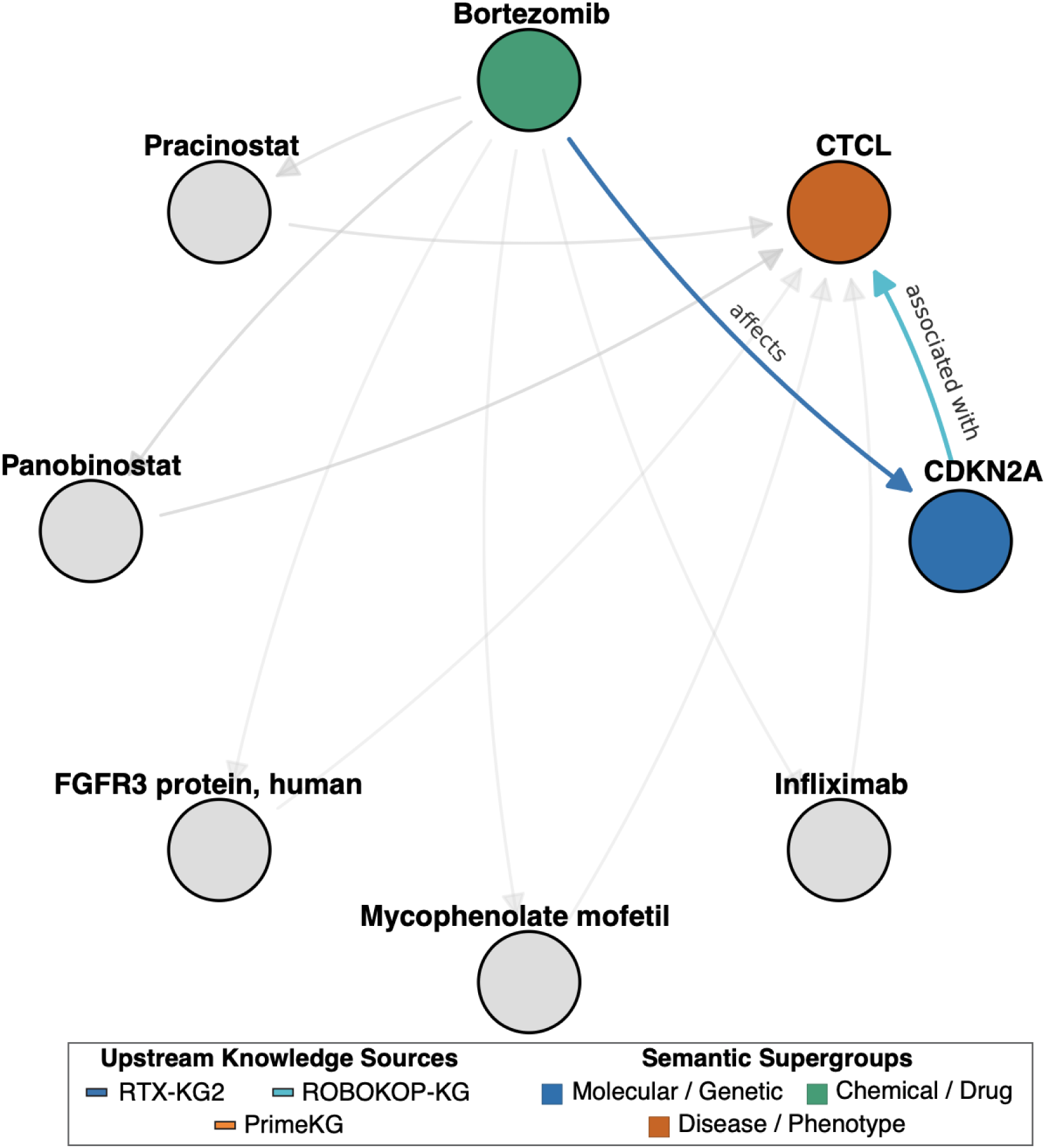
Subgraphs showing 2-hop SOPs across EC-KG for Bortezomib-Cutaneous T-Cell Lymphoma. Through integration of upstream networks, a novel pathway supporting mechanistic understanding of CTCL can be discovered.

### Prediction of Drug Repurposing Candidates

The integration of different networks into EC-KG not only provides explicit signals in the form of novel pathways, but also new patterns for improved ML training. Through the integration of different knowledge sources, signal in the form of disease-disease similarity can aid the process of identifying therapeutic candidates, with some ontology-guided machine learning models outperforming zero-shot foundation models.^15,74^ Therefore, we validated EC-KG’s utility for computational drug repurposing through an ML-based validation.

We trained random forest models to predict drug-disease treatment relationships. For this purpose, we used a subset of 40,818 drug-disease pairs present across all four networks, including 13,412 drug disease pairs with a known treatment relationship (y=1) and 27,406 contraindicated drug-disease pairs (y=0). For each KG, nodes were embedded with Node2Vec, and the drug and disease embeddings for a pair were concatenated to form its feature vector. Random Forest classifiers were then trained and evaluated using 5-fold cross-validation. To prevent leakage, all known treatment edges for evaluated drug-disease pairs were removed from each KG before embeddings. Models were evaluated on a withheld standard test set and on the curated off-label indication set, with any pair seen in training excluded from evaluation. To reflect the computational pharmacophenomics use case,^30^ we then generated treatment probabilities scores for the full cartesian product of the curated drug and disease lists.

Across both the standard test set and the out-of-distribution off-label set, EC-KG achieved the highest classification F1, significantly exceeding all three upstream KGs (Holm-adjusted p < 0.01; Fig. 8a). Because classification F1 at a fixed threshold is a summary, we additionally examined disease-specific Hit@k, which reveals distinct ranking profiles across the models (Fig. 8b, 8c). On the standard test set, models trained on PrimeKG showed the strongest disease-specific ranking (Fig. 8b), consistent with the curated nature of PrimeKG improving performance when identifying already-approved treatment relationships. On the off-label set, however, EC-KG achieved significantly higher ranking performance than PrimeKG, ROBOKOP KG, and RTX-KG2, as measured by area under the disease-specific Hit@1–100 curve (Holm-adjusted p ≤ 0.003); the corresponding Hit@10 advantage was significant against PrimeKG and ROBOKOP KG but not against RTX-KG2 (Fig. 8c).

**Figure 8.**
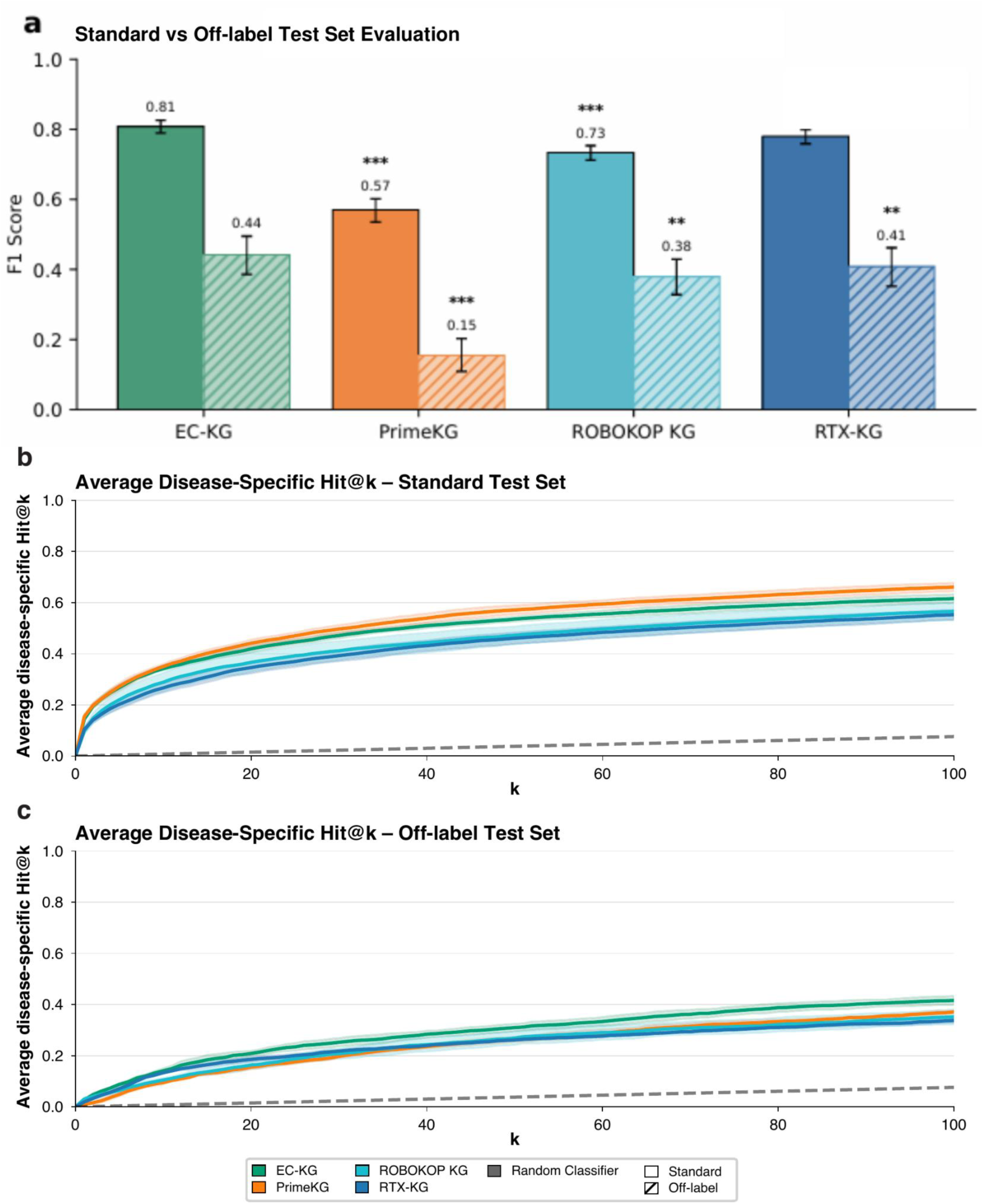
Performance of Random Forest classifiers trained on EC-KG and each upstream KG. a) Classification performance (F1 score) on the standard withheld test set (solid bars) and off-label evaluation sets (hatched bars), coloured by KG. Bars show the disease-level F1 pooled across the five held-out cross-validation folds (standard: test set includes 14,540 contraindications and 8,453 indications; off-label: the same contraindications and 612 off-label treatment pairs from the curated indications list). Error bars are 95% confidence intervals from a disease-clustered bootstrap (20,000 resamples). Asterisks denote the Holm-adjusted significance of the paired EC-KG-versus-comparator difference (two-sided disease-clustered permutation test; *** *p < 0.001, ** p < 0.01, * p < 0.05;* significance reflects the paired test, not the overlap of the per-bar intervals. b) Disease-specific ranking performance on the standard withheld test set (1706 on-label pairs across 872 diseases). c) Disease-specific ranking performance on off-label evaluation sets (612 off label indication pairs across 316 diseases). For all subplots in the figure, any pairs used during training were excluded from the evaluation set for all subplots. Statistically significant differences were found for AUC(Hit@1-100) across all and Hit@10 for EC-KG versus PrimeKG and ROBOKOP KG but not RTX KG2. Shaded bands in panels b-c indicate ±1 standard deviation across the five cross-validation folds.

We have also used the results of these predictions in the real world to identify potential repurposed treatments to the Every Cure medical team for further investigation, surfacing Botulinum Toxin A as a candidate to treat Major Depressive Disorder as well as to validate repurposing of Lenalidomide and Dexamethasone for a subgroup of patients with Rosai-Dorfman Disease.^75, 76^ This demonstrates how EC-KG and its unification of complementary biomedical networks enriches the biomedical signal within the latent space and can be beneficial for computational drug repurposing.

### Limitations

EC-KG and its construction pipeline have certain limitations. The normalization process and its coverage is restricted to the databases and ontologies already present in the Babel database. Therefore, integration of novel ontologies or custom data enrichment is challenging and requires manual mapping of identifiers. While the unification of different biomedical networks leads to emergence of novel context, it also has negative consequences on the topology of the graph, such as an increase in the number of ontologically or biomedically irrelevant pathways due to the combinatorial nature of network integration. Such a unification strategy can also lead to the presence of conflicting information within the graph: when two primary knowledge sources make different claims about the same subject-object pair, both edges will appear in EC-KG. We believe that through multi-level provenance tracking and knowledge-level attributes, users can mitigate noise and conflicting information in the graph: however, it remains one of the inherent limitations of KG integration.

Lastly, we note that while these results demonstrate EC-KG’s predictive utility on evaluation datasets and role in prioritizing drug repurposing opportunities, we have found that strong performance on data science metrics does not always translate to real-world plausibility of predicted repurposing opportunities, likely due to the complexity of biological systems and pharmacology. We consider true validation of these opportunities to come through in depth scientific review and ultimately clinical evaluation and testing of top hits, which is currently underway.

In conclusion, though additional work is needed to further advance these data sources and models and to efficiently evaluate them in the laboratory and clinic, EC-KG demonstrates that independently constructed biomedical knowledge graphs can be unified into an interoperable, provenance-rich network that improves the coverage and contextual representation of drugs and diseases relevant to repurposing.

## Usage Notes

EC-KG can be ingested into Neo4j or loaded into memory with graph-native libraries on researchers’ local machines. The instructions on how to load EC-KG from Hugging Face repository are detailed at https://docs.dev.everycure.org/releases/public_data_releases.

## Data Availability

EC-KG release v0.15.19 and its license is publicly available on Hugging Face (nodes: https://huggingface.co/datasets/everycure/kg-nodes; edges: https://huggingface.co/datasets/everycure/kg-edges) where future releases will also be uploaded. EC-KG can be explored through an independent KG dashboard (https://data.dev.everycure.org/versions/v0.15.19/evidence). The complete list of primary knowledge sources within EC-KG and their respective licenses is available at https://docs.dev.everycure.org/releases/primary_knowledge_sources. While EC-KG is provided in Hugging Face repository, for scientific reproducibility we also provide the preprocessed PrimeKG, RTX-KG2, and ROBOKOP KG to reproduce integration workflow (https://console.cloud.google.com/storage/browser/data.dev.everycure.org/data/01_RAW/KGs).

The curated drug and disease lists, and indications lists used for validation and topological metrics are also available on Hugging Face (drug list: https://huggingface.co/datasets/everycure/drug-list; disease list: https://huggingface.co/datasets/everycure/disease-list, indications list: https://huggingface.co/datasets/everycure/indications-list). The disease list contains the entire Mondo ontology however only a subset of diseases at ‘clinically recognized’ level was used as a core entity. For preservation and reproducibility, all of the datasets associated with the publication were archived at Zenodo with a DOI identifier (https://doi.org/10.5281/zenodo.20815442).

## Code Availability

The code for data preprocessing, integration, modelling, and infrastructure setup, and documentation, is available in Github repository at https://github.com/everycure-org/matrix. The documentation for navigating and working with the repository is available at https://docs.dev.everycure.org/. The code for figure generation or additional analysis for technical validation can be found in a separate repository at https://github.com/everycure-org/ec-kg.

